# mitoXplorer, a visual data mining platform to systematically analyze and visualize mitochondrial expression dynamics and mutations

**DOI:** 10.1101/641423

**Authors:** Annie Yim, Prasanna Koti, Adrien Bonnard, Milena Duerrbaum, Cecilia Mueller, Jose Villaveces, Salma Gamal, Giovanni Cardone, Fabiana Perocchi, Zuzana Storchova, Bianca H. Habermann

## Abstract

**Background:** Mitochondria produce cellular energy in the form of ATP and are involved in various metabolic and signaling processes. However, the cellular requirements for mitochondria are different depending on cell type, cell state or organism. Information on the expression dynamics of genes with mitochondrial functions (mito-genes) is embedded in publicly available transcriptomic or proteomic studies and the variety of available datasets enables us to study the expression dynamics of mito-genes in many different cell types, conditions and organisms. Yet, we lack an easy way of extracting these data for gene groups such as mito-genes.

**Results:** Here, we introduce the web-based visual data mining platform mitoXplorer, which systematically integrates expression and mutation data of mito-genes. The central part of mitoXplorer is a manually curated mitochondrial interactome containing ∼1200 genes, which we have annotated in 35 different mitochondrial processes. This mitochondrial interactome can be integrated with publicly available transcriptomic, proteomic or mutation data in a user-centric manner. A set of analysis and visualization tools allows the mining and exploration of mitochondrial expression dynamics and mutations across various datasets from different organisms and to quantify the adaptation of mitochondrial dynamics to different conditions. We apply mitoXplorer to quantify expression changes of mito-genes of a set of aneuploid cell lines that carry an extra copy of chromosome 21. mitoXplorer uncovers remarkable differences in the regulation of the mitochondrial transcriptome and proteome due to the dysregulation of the mitochondrial ribosome in retinal pigment epithelial trisomy 21 cells which results in severe defects in oxidative phosphorylation.

**Conclusions:** We demonstrate the power of the visual data mining platform mitoXplorer to explore expression data in a focused and detailed way to uncover underlying potential mechanisms for further experimental studies. We validate the hypothesis-creating power of mitoXplorer by testing predicted phenotypes in trisomy 21 model systems. MitoXplorer is freely available at http://mitoxplorer.ibdm.univ-mrs.fr. MitoXplorer does not require installation nor programming knowledge and is web-based. Therefore, mitoXplorer is accessible to a wide audience of experimental experts studying mitochondrial dynamics.

## Background

Enormous amounts of transcriptomic data are publicly available for exploration. This richness of data gives us the unique opportunity to explore the behavior of individual genes or groups of genes within a vast variety of different cell types, developmental or disease conditions or in different species. By integrating these data in a sophisticated way, we may be capable to discover new dependencies between genes or processes. Specific databases are available for mining and exploring disease-associated data, such as The Cancer Genome Atlas (TCGA [1]), or the International Cancer Consortium Data Portal (ICGC [2]). Especially cancer data portals provide users with the opportunity to perform deeper exploration of expression changes of individual genes or gene groups in different tumor types ([1–3]; for a review on available cancer data portals, see [4]). Expression Atlas on the other hand provides pre-processed data from a large variety of different studies in numerous species [5]. Indeed, the majority of transcriptomic datasets are not related to cancer and are stored in public repositories such as Gene Expression Omnibus (GEO [6]), DDBJ Omics Archive [7] or ArrayExpress [8]. Currently, it is not straightforward to integrate data from these repositories without at least basic programming knowledge.

Next to extracting reliable information from -omics datasets, it is equally important to support interactive data visualization. This is a key element for a user-guided exploration and interpretation of complex data, facilitating the generation of biologically relevant hypotheses – a process referred to as visual data mining (VDM, reviewed e.g. in [9]). Therefore, essentially all online data portals provide graphical tools for data exploration.

What is fundamentally lacking is a user-centric, web-based and interactive platform for data integration of a set of selected genes or proteins sharing the same cellular function(s). The benefits of such a tool are evident: first, it would give us the possibility to explore the expression dynamics of or mutations in this set of selected genes across many different conditions, tissues, as well as across different species. Second, by integrating data using enrichment techniques, for instance with epigenetic or ChIP-seq data or by network analysis using the cellular interactome(s), we can recognize the mechanisms that regulate the expression dynamics of the selected gene set.

One interesting set of genes are mitochondria-associated genes (mito-genes): in other words, all genes, whose encoded proteins localize to mitochondria and fulfill their cellular function within this organelle. Mito-genes are well-suited for such a systematic analysis, because we have a relatively complete knowledge of their identity and can categorize them according to their mitochondrial functions [10]. This *a priori* knowledge can help us in mining and exploring the expression dynamics of mito-genes and functions in various conditions and species.

Mitochondria are essential organelles in eukaryotic cells that are required for producing cellular energy in form of ATP and for numerous other metabolic and signaling functions [10]. Attributable to their central cellular role, mitochondrial dysfunctions were found to be associated with a number of human diseases such as obesity, diabetes, neurodegenerative diseases and cancer [11–15]. However, mitochondria are not uniform organelles. Their structural and metabolic diversity and how they influence each other has been well described in literature [16–20]. This mitochondrial heterogeneity in different tissues is reflected in their molecular composition [21]. The total number of proteins that contribute to mitochondrial functions and localize to mitochondria is currently not precisely known and might differ between tissues and species [22,23]. Yet, based on proteomics data from several organisms, it is likely that mitochondria contain more than 1000 proteins [23–30]. Mitochondria have their own genome, whose size in animals is between 11 and 28 kilo-bases [31]. Most metazoan mitochondria encode 13 essential proteins of the respiratory chain required for oxidative phosphorylation (OXPHOS), all rRNAs of the small and large mitochondrial ribosomal subunits, as well as most mitochondrial tRNAs [32]. All other proteins found in mitochondria (mito-proteins) are encoded by genes in the nucleus; the protein products of these nuclear-encoded mitochondrial genes (NEMGs) are transported to and imported into mitochondria.

Based on data from mitochondrial proteomics studies or genome-scale prediction of mito-proteins, several electronic repositories of the mitochondrial interactome have been created [24,33–36], though they often lack proper functional assignments of mito-proteins. Moreover, proteomics studies describing the mitochondrial proteome can suffer from a high false-positive rate [23]. Computational prediction or machine learning [37] on the other hand lack experimental confirmation. As a consequence, none of the published mitochondrial interactomes available to date can be taken without further manual curation. Moreover, these lists are not integrated with any available data analysis tool to explore mitochondrial expression dynamics under varying conditions or in different tissues or species.

In this study, we present mitoXplorer, a web-based, highly interactive visual data mining platform to integrate transcriptome, proteome, as well as mutation-based data with a manually curated, function-based mitochondrial interactome. With mitoXplorer, we can explore the expression dynamics, as well as mutations of mito-genes and their associated mitochondrial processes (mito-processes) across a large variety of different -omics datasets without the need of programming knowledge. MitoXplorer provides users with dynamic and interactive figures, which instantly display information on mitochondrial gene functions and protein-protein interactions. To achieve this, mitoXplorer integrates publicly available -omics data with our hand-curated mitochondrial interactomes for different model species. Additionally, users can upload their own data for integration with our hand-curated mitochondrial interactome, as well as the publicly available -omics data stored in the mitoXplorer database. In order to test the analytical and predictive power of mitoXplorer, we generated transcriptome and proteome data from aneuploid cell lines, carrying trisomy 21 (T21). We used mitoXplorer to analyze and integrate our data with publicly available trisomy 21 data. MitoXplorer enabled us to predict respiratory failure in one of our T21 cell lines, which we experimentally confirmed, demonstrating the predictive power of mitoXplorer.

## Results

The outline of the mitoXplorer web-platform is illustrated in Figure 1: at the *back-end*, manually curated mitochondrial interactomes from human, mouse and Drosophila, as well as expression and mutation data from these three species are stored in a MySQL database (details on the implementation of the *back-end* are available in Methods, as well as Additional File 1, Supplementary Figure S1).

**Figure 1:**
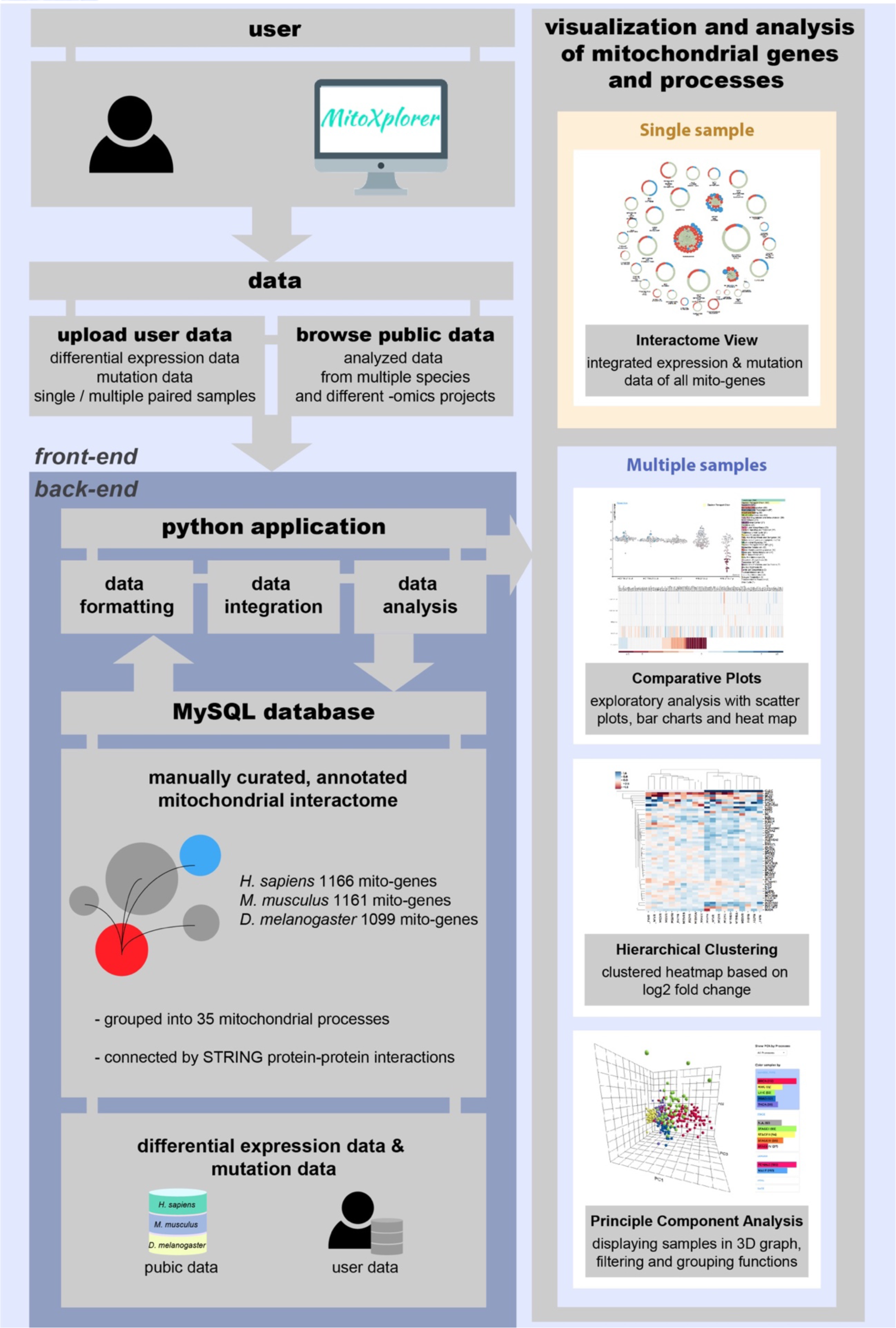
Setup of the mitoXplorer web-based visual data mining platform. A manually curated, annotated mitochondrial interactome represents the central part of the mitoXplorer software, for which we have assembled 1166 mito-genes in human, 1161 mito-genes in mouse and 1099 mito-genes in fruit fly in 35 mitochondrial processes (mito-processes). We have connected gene products using protein-protein interactions from STRING [39]. Publicly available expression and mutation data from repositories such as TCGA or GEO are provided for data integration, analysis and visualization and are stored together with species interactomes in a MySQL database. Users can provide their own data, which are temporarily stored and only accessible to the user. A set of Python-based scripts at the *back-end* of the platform handle data formatting, integration and analysis (Additional File 1, Supplementary Figure S1). The user interacts with mitoXplorer via several visual interfaces, by which the user can analyze, integrate and visualize his private, as well as public data. Four interactive visualization interfaces are offered: 1) the Interactome View allows at-a-glance visualization of the entire mitochondrial interactome of a single dataset (see Figure 2); 2) Comparative Plots, consisting of a scatter plot and a sort-able heatmap allows comparison of up to six datasets, whereby a single mito-process is analyzed at a time (see Figure 3); 3) Hierarchical Clustering allows comparison of a large number of datasets, whereby datasets are clustered according to their expression values. Hierarchical Clustering plots are zoom-able and interactive (see Figure 4); 4) Principle Component Analysis displays PCA-analyzed datasets in 3D, providing filtering and grouping functions. There is in principle no limit to the number of datasets that can be analyzed using PCA (see Figure 5).

The user interacts with the mitoXplorer web-platform via the *front-end*, which offers different visualization and analysis methods. Users can either browse stored public data or upload their own data.

### The mitochondrial interactomes

The main component of mitoXplorer is the mitochondrial interactome. Its accurate annotation and completeness are essential for performing a meaningful mitoXplorer-based analysis. To establish mitochondrial interactomes, we have assembled and manually curated lists of genes with annotated mitochondrial processes (mito-processes) for human, mouse and Drosophila. Starting from published mitochondrial proteomics data [27], we removed false-positives and supplemented likely missing genes using information from Mitocarta [24], as well as orthologs across species. We relied mainly on literature sources and information from the respective gene entry at NCBI [38] for establishing whether a protein in question is primarily localized to mitochondria. This resulted in 1166 human, 1161 mouse and 1099 Drosophila mito-genes. We annotated the genes with, and grouped them according to 35 mito-processes using controlled vocabulary (Table 1). In addition to purely mitochondrial processes, we added cytosolic processes coupled to mitochondrial functions, including glycolysis, the pentose phosphate pathway or apoptosis. According to our annotation strategy, one gene is part of only a single mito-process. We used all mito-genes to create mitochondrial interactomes by adding protein-protein interaction information from STRING [39].

**Table 1:** Mito-processes and number of genes in Human, Mouse and Drosophila.

| Mito-process | Human | Mouse | Drosophila |
| --- | --- | --- | --- |
| Amino Acid Metabolism | 81 | 80 | 67 |
| Apoptosis | 55 | 54 | 44 |
| Bile Acid Synthesis | 2 | 2 | 7 |
| Calcium Signaling and Transport | 24 | 23 | 12 |
| Cardiolipin Biosynthesis | 5 | 5 | 5 |
| Fatty Acid Biosynthesis & Elongation | 22 | 22 | 15 |
| Fatty Acid Degradation & Beta-oxidation | 30 | 31 | 26 |
| Fatty Acid Metabolism | 14 | 11 | 19 |
| Fe-S cluster biosynthesis | 24 | 25 | 18 |
| Folate & Pterine Metabolism | 12 | 12 | 9 |
| Fructose Metabolism | 7 | 7 | 3 |
| Glycolysis | 37 | 39 | 35 |
| Heme Biosynthesis | 9 | 9 | 9 |
| Import & Sorting | 51 | 52 | 62 |
| Lipoic Acid Metabolism | 3 | 3 | 4 |
| Metabolism of Lipids & Lipoproteins | 34 | 36 | 17 |
| Metabolism of Vitamins & Co-Factors | 16 | 15 | 18 |
| Mitochondrial Carrier | 46 | 45 | 46 |
| Mitochondrial Dynamics | 60 | 60 | 47 |
| Mitochondrial Signaling | 18 | 18 | 10 |
| Nitrogen Metabolism | 8 | 8 | 21 |
| Nucleotide Metabolism | 14 | 14 | 13 |
| Oxidative Phosphorylation | 167 | 164 | 174 |
| Oxidative Phosphorylation (MT) | 13 | 13 | 13 |
| Pentose Phosphate Pathway | 7 | 7 | 6 |
| Protein Stability & Degradation | 26 | 26 | 20 |
| Pyruvate Metabolism | 26 | 25 | 24 |
| Replication & Transcription | 53 | 54 | 33 |
| **ROS Defense** | 32 | 32 | 30 |
| **Translation** | 184 | 183 | 191 |
| **Translation (MT)** | 24 | 24 | 24 |
| **Transmembrane Transport** | 20 | 19 | 21 |
| **Tricarboxylic Acid Cycle** | 21 | 22 | 29 |
| **Ubiquinone Biosynthesis** | 9 | 9 | 9 |
| **Unknown** | 13 | 13 | 21 |

Currently, the interactomes of three organisms are available on mitoXplorer: *Homo sapiens* (human), *Mus musculus* (mouse) and *Drosophila melanogaster* (fruit fly). Mito-genes of human, mouse and Drosophila annotated with mito-processes are available in Additional File 2, Supplementary Table S1 a-c. These manually curated and annotated interactomes enables the meaningful analysis and visualization of mitochondrial expression dynamics of mito-processes by comparing differential expression of two or more conditions in mitoXplorer.

### The mitoXplorer expression and mutation database

To foster the analysis of mitochondrial expression dynamics and mutations, mitoXplorer hosts expression and mutation data from public repositories in a MySQL database.

Expression data encompass analyzed data of differentially expressed genes from RNA-seq studies and are available in the form of log2 fold change (log2FC) and p-value. One differential dataset thus includes two experimental conditions with all replicates. Mutation data include analyzed data of identified SNPs of one sample against a publicly available reference genome or transcriptome. Pre-analyzed public data are taken as provided by the authors of the respective study. Thus, the algorithms and their settings might differ between data from different studies or sources.

The largest public resource imported into mitoXplorer covers publicly available expression data of human cancers from The Cancer Genome Atlas (TCGA, [1]). We have included all paired samples. This resulted in a total of 523 differential datasets from 6 different cancer types: kidney cancer (KIRK), breast cancer (BRCA), liver cancer (LIHC), thyroid cancer (THCA), lung cancer (LUAD) and prostate cancer (PRAD). Changes in mitochondrial metabolism have been described in many cancer types (for a review, see [40]). As mitoXplorer is the thus far only resource that allows a focused analysis of mito-genes across different cancer types or patient groups, this resource should be especially useful to shed light on the expression dynamics or mutational data of mito-genes in cancer and to classify the mitochondrial metabolic profiles of tumor types and sub-types. Users can moreover integrate proprietary data with differential expression or with mutation data from different tumor types and subtypes.

We provide data from human trisomy 21 patients (GEO accession numbers: GSE55426; GSE79842; [41,42]), from trisomy 21 studies in mouse (GSE5542 [41]; GSE79842 [42]; GSE48555 [43]), as well as differential datasets from this study from human trisomic cell lines (11 datasets), which have been partially published elsewhere [44,45] (GEO accessions: GSE39768; GSE47830; GSE102855). These transcriptomic, as well as proteomic datasets should help understand the role of mitochondria and the mitochondrial metabolism in trisomy 21.

We also uploaded differential transcriptomic and proteomic data of five different mouse conditional heart knock-out strains of genes involved in mitochondrial replication, transcription and translation [46] (Lrpprc, Mterf4, Tfam, Polrmt, Twnk (Twinkle), (GEO accession: GSE96518)). These data are especially helpful in unraveling the transcriptional and post-transcriptional effects on mito-genes upon disruption of gene expression at different levels in mitochondria.

To extend mitoXplorer to other model organism, we added data from *D. melanogaster*, namely expression data from 185 wild-derived, inbred strains (males and females) from the Drosophila Genetics Reference Panel [47] (DGRP2). The wild-derived fly strains come from different environmental and social situations and display a substantial quantitative genetic variation in gene expression. The availability of these data on mitoXplorer allows a focused analysis of mito-genes to elucidate, whether mitochondrial expression dynamics is equally impacted by the environment in these strains. Finally, we have uploaded data from a recently published systematic study of flight muscle development in *D. melanogaster* [48] (GEO accession: GSE107247). This enables the analysis of mitochondrial expression dynamics during the development and differentiation of a tissue that is highly dependent on an efficient mitochondrial metabolism and especially ATP production for proper functioning.

All publicly available data can be viewed and accessed from the mitoXplorer DATABASE web-site.

### User-provided expression and/or mutation data

Researchers can upload and explore their own data in mitoXplorer, given that they originate from one of the species contained in the mitoXplorer platform. Data must be pre-analyzed. Differential expression data must contain the dataset ID (describing the experimental condition), the gene name and the log2FC. Optional values include the p-value, as well as the averaged read counts (or intensities) of the replicates of the compared conditions. Mutation data must contain the dataset ID, gene name, the chromosome, the position, as well as reference and alternative allele. Optional values include the effect, as well as the consequence of the mutation. The entire list of genes from a study should be uploaded to the platform for several reasons: first, a restriction to only differentially expressed or mutated genes will suppress links between proteins in the interactome; second, an integration of user data with publicly provided data is difficult with incomplete datasets; third, mitoXplorer will automatically select the mito-genes from the user data. Users can either generate their own data in the format described on our website; or use the RNA-seq pipeline that we provide at https://gitlab.com/habermannlab/mitox_rnaseq_pipeline/. Uploaded data will be checked for correct formatting and integrated with the interactome of the chosen species. User data are only visible to the owner and are stored in the mitoXplorer MySQL database for 7 days. Users can integrate their own data with available public data on mitoXplorer to perform various analyses and visualizations as described below (Figure 1).

### Analysis and visualization tools in mitoXplorer

The mitoXplorer web-platform provides a set of powerful, easy-to-read and highly interactive visualization tools to analyze and visualize public, as well as user-provided data by VDM (Figure 1): an **Interactome View** to analyze the overall expression and mutation dynamics of all mito-processes of a single dataset containing differentially expressed genes between two conditions and potential mutations in mito-genes; the **Comparative Plot**, consisting of an interactive scatter plot, as well as an interactive heatmap for comparing up to 6 datasets; the **Hierarchical Clustering**, as well as the **Principal Component Analysis** for comparative analysis of many datasets.

## Interactome View

The Interactome View can be used to get an at-a-glance view of the overall expression dynamics of all mito-processes of a single dataset of differentially expressed mito-genes and potential mutations (Figure 2 a). It allows users to identify the most prominently changed mito-processes or -genes in a dataset. The genes are grouped according to mito-processes and displayed in the process they are assigned to. The Interactome View is highly dynamic and can be adjusted by users to their needs.

**Figure 2:**
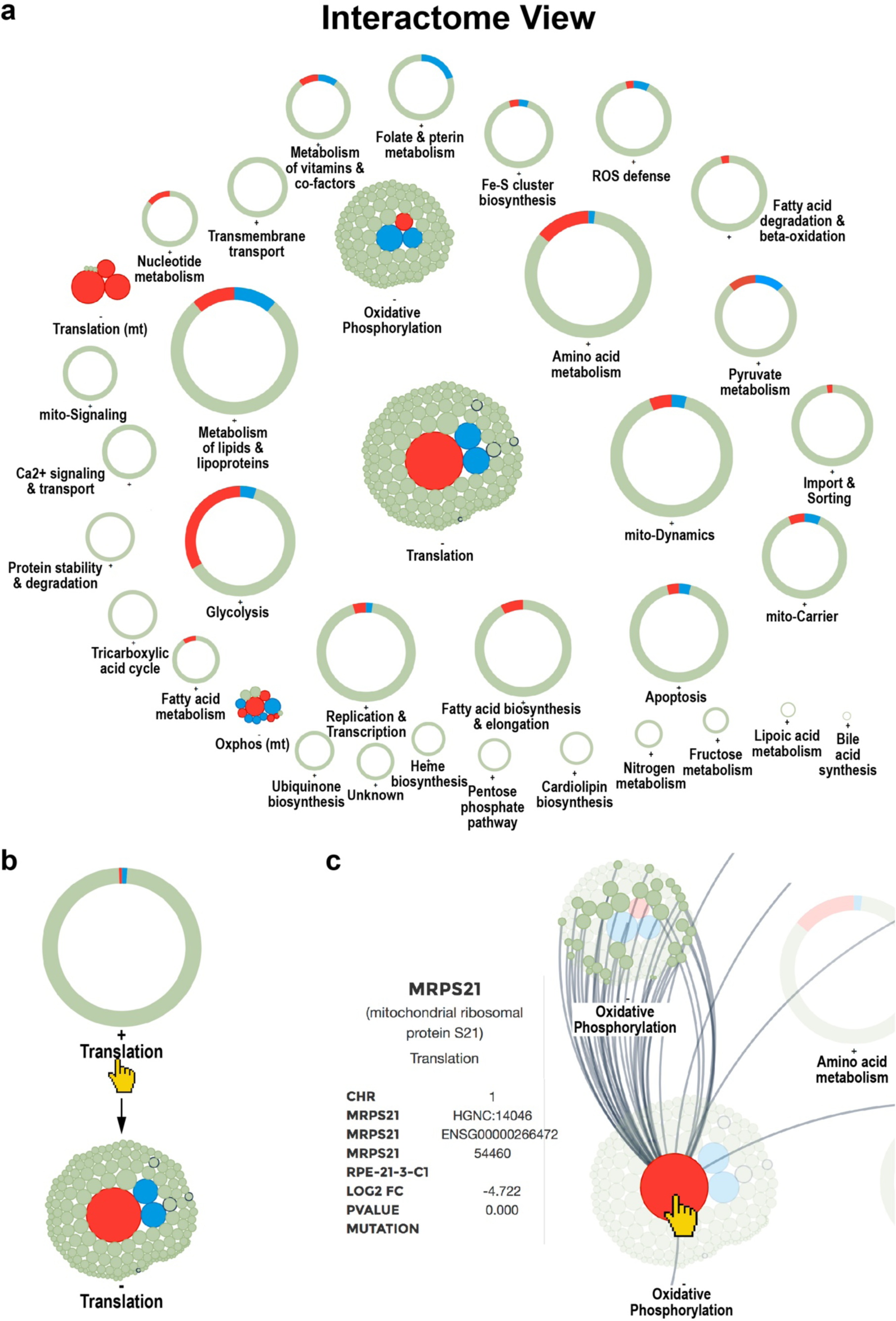
Interactome View of the mitoXplorer platform. **(a)** Overview of all mito-processes of one dataset. A process can either be shown as one circle with colored segments according to the number of dysregulated genes, or upon clicking on the process, by showing all individual genes being part of this process; see **(b)** by clicking on the process Translation or the adjacent ‘+’, the circle is replaced by individual bubbles representing genes of this process. Clicking on the process again, or on the adjacent ‘-’ will revert to the circular display. **(c)** Hovering over a gene bubble will display the name of the gene and associated information (gene name, description, chromosomal location, mitochondrial process, accession numbers, as well as log2 fold change, p-value and observed mutations), as well as all connections to mito-genes in other processes. Compared were the retinal epithelial cell line RPE1 (RPE) wild-type to RPE1 with Trisomy 21 (RPE_T21).

When the Interactome View is launched, each mito-process is primarily shown as a grey circle with elements colored in grey, blue and/or red, indicating up- or down-regulated genes within the process, respectively (Figure 2 a). Thus, mito-processes with the most up- or down-regulated genes can be quickly identified.

When clicking on a process name, its circle opens and displays all its member genes as bubbles, whereby the size of the bubble indicates the strength of the differential regulation and the color indicates up- (blue) or down- (red) regulation of the gene (Figure 2 b). Both, the log2FC as well as the p-value are color-coded in the Interactome View. Only genes with a p-value below 0.05 will be colored. If information about mutations are included in the dataset, these are indicated by a thicker, black border of the gene bubble.

Hovering over a gene will display the gene name, its function, its mito-process, the log2FC and the p-value of the differential expression analysis, as well as potential mutations in the *information panel* (Figure 2 c). If a gene physically interacts with other mito-genes, hovering over it or over the process circle will in addition display these connections (Figure 2 c). Thus, the user is immediately informed about the location and connectivity of the protein of interest within the mitochondrial interactome. Users can also search for specific genes using the ‘FIND A GENE’ box at the top of the page.

The Interactome View can be launched by clicking on the ‘eye’ symbol next to dataset names from the ANALYSIS page of mitoXplorer, after having chosen the organism, the project and the dataset. Alternatively, users can access single datasets from the DATABASE page of the platform, by clicking on the eye symbol of a listed dataset after having chosen a species, as well as a project. A new page will be opened for the Interactome View, which allows opening and comparing multiple datasets at the same time. This is especially useful for comparing the overall expression change of mito-processes of multiple datasets.

## Comparative Plot

The Comparative Plot visualization combines several interactive graphs to analyze one mito-process, allowing the comparison of up to 6 datasets. It includes a scatter plot with a dynamic y-axis, as well as an interactive heatmap at the bottom of the page. The mito-process to be visualized can be selected in the *process panel* (Figure 3 a). Red and blue coloring of the dots and the bar chart indicates the directionality of differential expression (blue: upregulated; red: downregulated); bright blue, larger gene bubbles in the scatter plot indicate mutations, if available from the dataset. This view offers an overview of the expression dynamics of all members of one mito-process for up to 6 individual datasets and thus can be helpful in identifying co-regulated genes e.g. in time-course data, patients or multiple mutant datasets.

**Figure 3:**
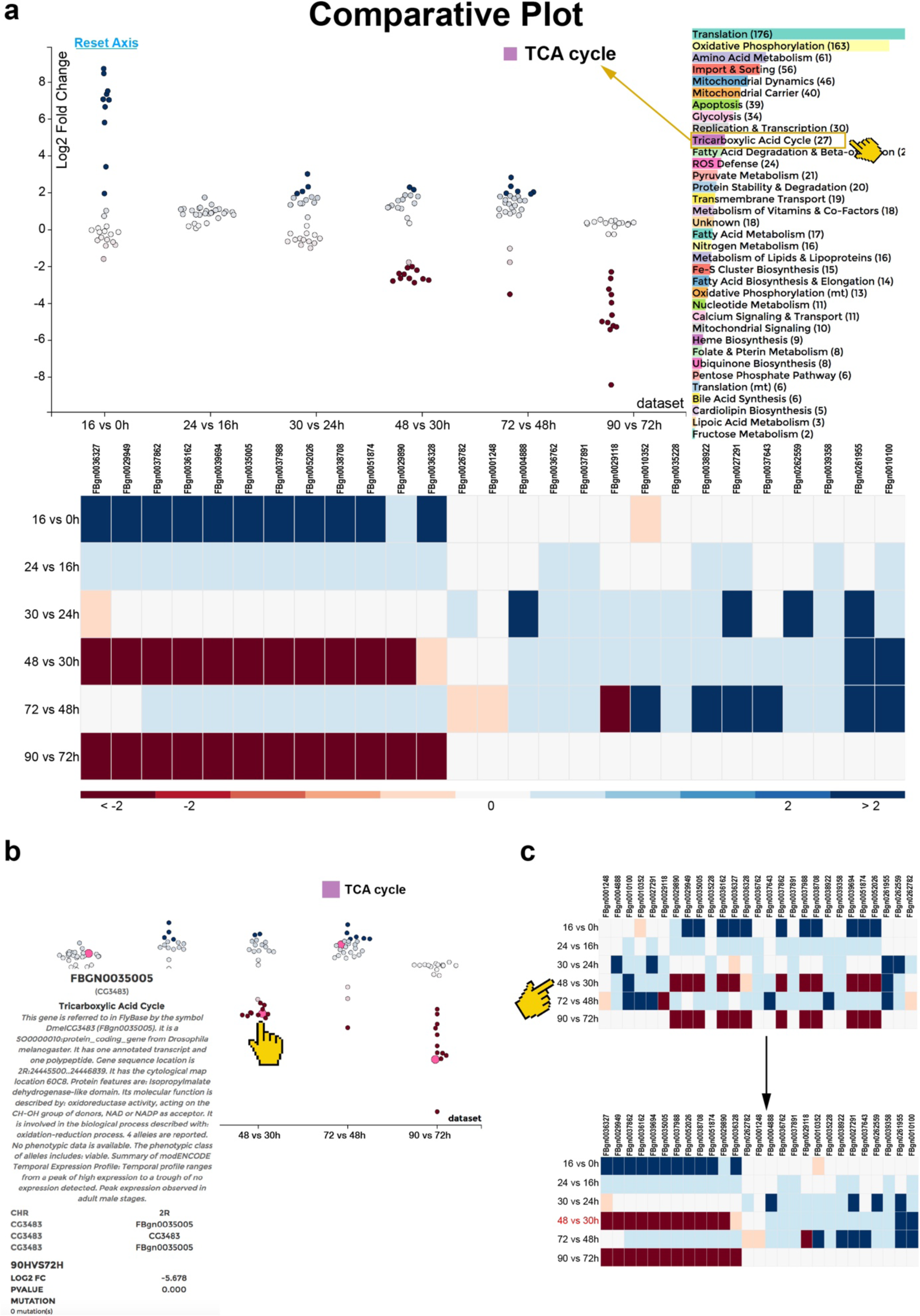
Comparative Plot of the mitoXplorer platform. **(a)** The Comparative Plot display is composed of a scatterplot and sortable heatmap and a bar chart for the selection of mito-processes. The scatterplot shows the log2 fold change (y-axis) and the datasets (x-axis). Each bubble represents one gene, whereby red dots indicate downregulated, and blue dots upregulated genes. The process to be shown can be selected by clicking on the process name in the bar chart next to the scatterplot, the chosen process is indicated on its top. In this case, TCA cycle was chosen. The heatmap at the bottom shows the individual genes and the datasets, whereby the genes are colored according to their log2 fold change (indicated at the bottom of the plot). **(b)** Hovering over a gene bubble (or a gene square in the heatmap) will display available information (in case of fly: gene name, mitochondrial process, gene description, chromosomal location, gene symbol, as well as log2 fold change, p-value and observed mutations). **(c)** The heatmap is sortable by log2 fold change (as indicated by the pointer in **c**), as well as by dataset. Clicking on one of the datasets will sort the heatmap according to the log2 fold change of all genes in this dataset, as is illustrated here. Clicking on one of the genes will sort the heatmap according to its log2 fold changes across different datasets. The time-series study of developing flight muscle was used to demonstrate the functionality of this visualization method.

Hovering over a gene bubble, or over a bar in the heatmap will again display the respective associated information of the gene in the *information panel* (gene name, function, mito-process, log2FC, p-value, potential mutations) (Figure 3 b). The heatmap can be sorted according to the dataset, as well as the differential expression values within one dataset (Figure 3 c). The Comparative Plot is especially useful for performing a detailed, comparative, mito-process based analysis of differential expression dynamics between different datasets.

We applied this analysis method to visualize differential expression data from a time-series study of flight muscle development during pupal stages in Drosophila [48] (Figure 3). While enrichment analysis has revealed a general positive enrichment of processes like ‘TCA Cycle’ in the course of flight muscle development, mitoXplorer identifies 12 genes of TCA cycle that are co-regulated. This group of genes is strongly upregulated between 0 and 16 hours of development, when myoblasts divide and fuse to myotubes. The same group of genes is consecutively downregulated in two phases at time-points 30 to 48 hours and 72 to 90 hours APF, when myotubes differentiate to mature muscle fibers. This is surprising as in mature muscle fibers, the TCA cycle should be important for proper functioning. Their strong induction between the first two time-points could be responsible for downregulation at later stages.

## The Heatmap: Hierarchical Clustering

Hierarchical Clustering visualization allows the analysis of up to 100 datasets, analyzing one process at a time. This creates a heatmap with mito-genes, as well as -datasets, which are clustered according to the log2FC using hierarchical clustering (Figure 4 a). The results are displayed as a clustered heatmap, with a dendrogram indicating the distance between datasets or between genes.

**Figure 4:**
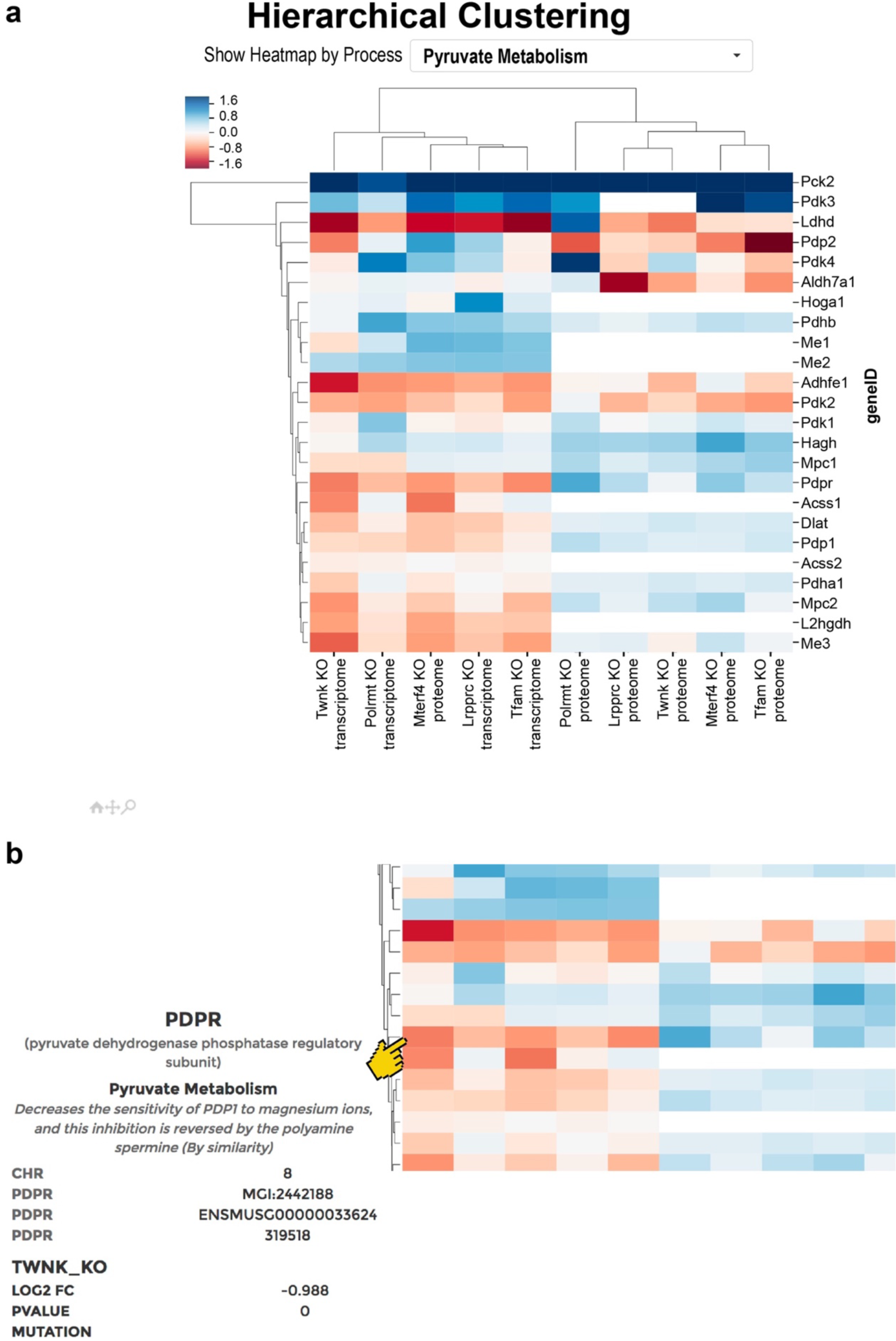
Hierarchical Clustering and heatmap plot of the mitoXplorer platform. Hierarchical Clustering of expression data results in a so-called heatmap. **(a)** Heatmap of transcriptome and proteome data of mouse knock-out strains of genes involved in mitochondrial replication, DNA-maintenance, transcription and RNA processing (taken from [46]). Data are clustered according to genes, as well as datasets. Gene boxes are colored according to their log2 fold change. At the top of the heatmap, the user can choose the mito-process to be displayed. **(b)** Hovering over one of the gene boxes will display information on the gene, such as the gene name, mito-process, log2 fold change, p-value and – if available – observed mutations. The heatmap is also zoom-able by clicking on the magnification glass at the bottom of the plot, so that large datasets can be visualized and analyzed efficiently. Datasets can be selected within the heatmap by grouping. To do this, first a group name has to be defined; second, the datasets belonging to this group have to be selected by clicking on one of the gene boxes of the dataset. This process can be repeated and the resulting groups can then be analyzed using Comparative Plots.

Hovering over a gene will display its associated information, as well as dataset information in the *information panel* (Figure 4 b). The user can furthermore zoom into parts of the heatmap to get a more detailed view of the data. The heatmap is particularly useful for discovering groups of similarly regulated mito-genes or datasets within one mito-process.

We applied this visualization tool to display transcriptome and proteome data from a recent, systematic study of mouse conditional knock-out strains for five genes involved in mitochondrial replication (*Twinkle (Twnk)*), mtDNA maintenance (*Tfam*), mito-transcription (*Polrmt*), mito-mRNA maturation (*Lrpprc*) and mito-translation (*mTerf4*) [46]. Interestingly, the expression dynamics of the mitochondrial transcriptomes and proteomes in heart tissue did not cluster together for the mutants upon the loss of any of these genes. In accordance with this, the expression of some mito-genes in the process pyruvate metabolism that is shown here differs on transcriptome and proteome level. This demonstrates the usefulness of hierarchical clustering and the heatmap display in identifying the correlation or divergence between genes as well as datasets.

## Principal Component Analysis

A larger number of datasets can be compared using Principal Component Analysis (PCA), either for an individual mito-process, or considering all mito-genes together (Figure 5 a). In PCA, the expression value (e.g. log2FC) of each gene is considered as one dimension, and each dataset represents one data point. In the resulting 3D PCA plot, the three axes represent the first three principal components and each bubble represents one dataset. The PCA is again interactive. The mito-process to be viewed can be selected via a drop-down menu on the top of the page. The plot can be turned and moved in 3D and has a zooming function.

**Figure 5:**
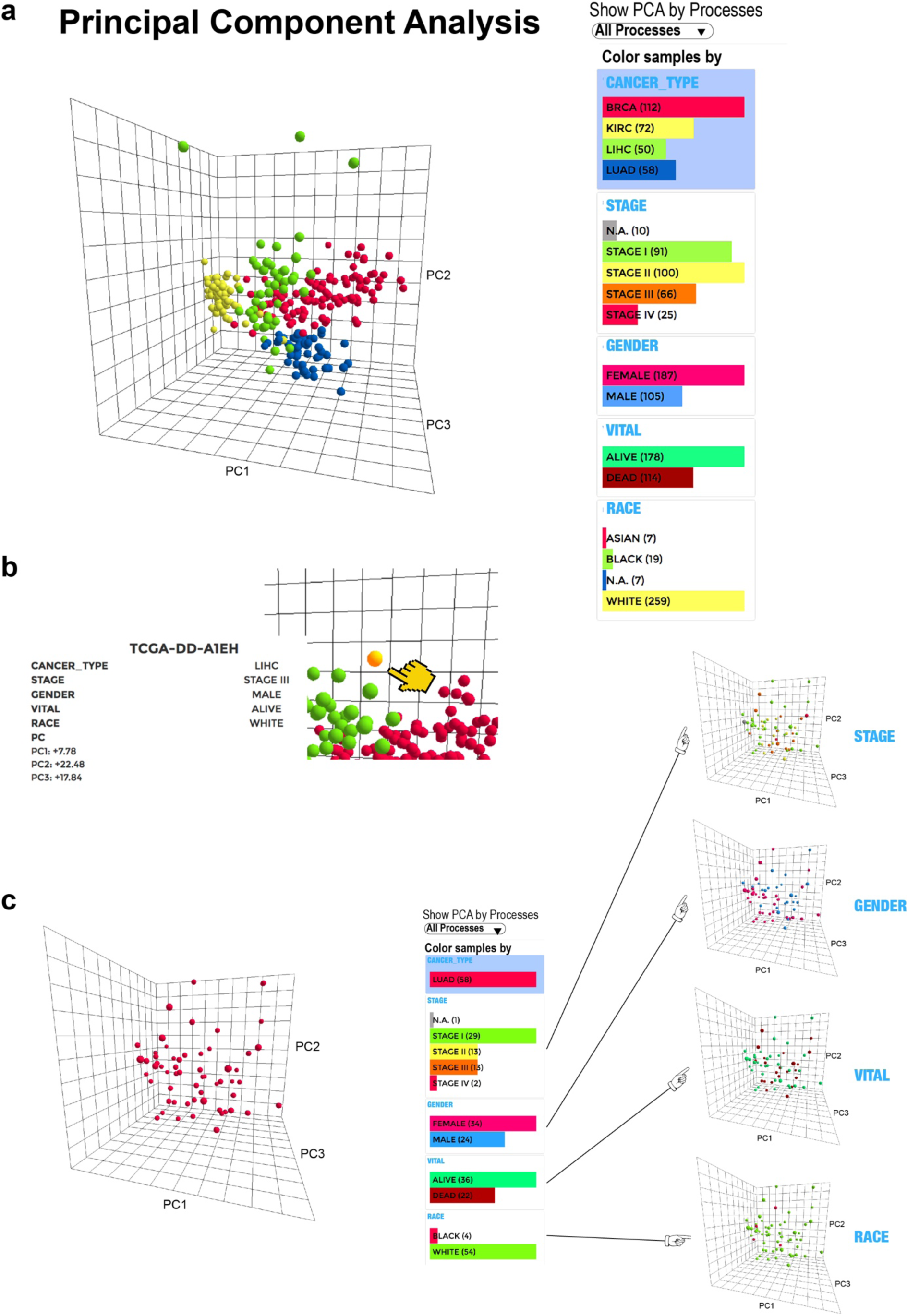
Principal component analysis and PCA plot of the mitoXplorer platform. **(a)** PCA analysis and plot of transcriptome data of The Cancer Genome Atlas (TCGA) database [1], showing four different cancer types: breast cancer (BRCA), kidney cancer (KIRK), liver cancer (LIHC) and lung cancer (LUAD). Each bubble represents one dataset, in this case, one cancer patient. At the right side at the top of the plot, the mito-process to be shown can be chosen. In this case, ‘All Processes’ are chosen, containing data from all mito-genes. At the right side next to the plot, different colors, as well as filters can be chosen. In this case, the Cancer Type was chosen for coloring, showing the four different cancer types in four different colors. **(b)** Hovering over a bubble will display associated information on the dataset, including the dataset name, and in case of the TCGA, information on the cancer type, the stage, the gender, the vital status, as well as skin color. In addition, the three PC components are shown. **(c)** Selecting color schemes on the right-hand side will change the coloring of the bubbles. In this case, only lung cancer is shown, and coloring is done according to Stage, Gender, Vital, and Skin color. This panel can also be used for selecting specific datasets. For instance, clicking on one of the stages will only display the chosen stage and omit datasets from other stages. As in the heatmap, datasets can be selected from the PCA for grouping. To do this, first a group name has to be defined; second, the datasets belonging to this group have to be selected by clicking on one of the dataset bubbles. This process can be repeated and the resulting groups can then be analyzed using Comparative Plots.

Hovering over a bubble will give all information associated with the individual dataset in the *information panel*, including the values of the first three principal components (Figure 5 b). The information differs for each project chosen.

Individual datasets can be selected and colored via the *dataset panel* next to the plot (Figure 5 c). For data from TCGA, the filter and coloring can for instance be used to highlight or to limit the plot to data from different tumors, different tumor stages or according to any other additional information provided. The PCA is especially useful for analyzing a large number of datasets and displaying specific trends in sub-groups.

We used the PCA plot to visualize data from the TCGA for four cancer types stored in mitoXplorer in Figure 5 a, whereby the colors of the bubbles represent the different tumor types. The PCA mode clearly highlights the distinctness of the different tumor types. In particular, kidney and liver cancer are highly distinct with respect to the first three components of all mito-genes (Figure 5 a).

Both views intended for large datasets, the Heatmap and the PCA, can identify groups of correlated datasets. In order to allow a more detailed, gene-centered analysis of correlated datasets, we added the possibility to select and group datasets in the Heatmap and the PCA view. Groups of datasets can be compared against each other with the Comparative Plot, whereby the log2FC is averaged over the data within a group. This functionality is useful, if for instance different groups of patients with a similar expression pattern need to be compared to each other; or to compare the expression changes during tumor development in different tumor stages.

Taken together, mitoXplorer provides a versatile, interactive and integrative set of tools to visualize and analyze the expression dynamics as well as mutations of mito-genes and mito-processes, providing a detailed understanding of observed changes at a molecular level.

### Analyzing cell lines carrying trisomy 21 using the mitoXplorer platform

To demonstrate the analytical and predictive power of mitoXplorer, we analyzed the transcriptome and proteome of a set of aneuploid cell lines carrying an extra copy of chromosome 21 (trisomy 21, T21). Mitochondrial dysfunction has been repeatedly found in T21 patients, whereby mostly oxidative stress, as well as – potentially resulting – mitochondrial respiratory deficiency have been shown to contribute to some of the observed clinical features (see for instance [49–62]). Transcriptome studies of different T21 tissues using microarrays [63–73] and more recently RNA sequencing [41,42,74] and proteomics [75–79] have revealed a complex picture of gene expression changes, with a marked dissimilarity in differential expression of mito-genes on mRNA and protein levels, indicating a potential post-transcriptional regulatory effect of some mito-genes in T21 [78]. Yet, mito-gene and protein expression data in different tissues or under varying conditions in T21 remain still sparse and a coherent hypothesis of the underlying mechanisms leading to the mitochondrial deficiencies in T21 patients is still missing.

We used trisomy 21 cell lines derived from either the euploid human colon cancer cell line HCT116 or from the retinal pigmented epithelial cell line RPE1, to which an extra copy of chromosome 21 [45] was added. We used two RPE1-derived and two HCT116-derived clones trisomic for chromosome 21 (Additional File 3, Supplementary Table S2 a), which were validated by fluorescent *in situ* hybridization and by whole genome sequencing. We used transcriptomic data of the original euploid RPE1 line and its two trisomic derivatives (RPE_T21 clone 1 and 2 (c1, c2) [45]), as well as for HCT116, and its trisomic derivatives (HCT_T21 (c1, c3)). We included proteomics data for RPE1 and one of its T21 derivatives (RPE_T21 c1). We performed bioinformatic analysis to determine differential expression of the above conditions (Additional File 3, Supplementary Table S2 b-e) and uploaded the differential expression data of the transcriptome and proteome on the mitoXplorer platform for further in-depth, mitochondrial analysis.

### Differences between trisomy 21 cell lines

MitoXplorer analysis of data comparing HCT116- and RPE1-derived T21 cell lines using the Interactome View revealed that T21 induced strong effects with respect to the overall expression changes in mito-genes (Figure 6). HCT_T21 showed a subtle, but consistent up-regulation of mito-genes (Figure 6 a). In contrast, RPE_T21 cells showed a strong down-regulation of a few genes involved in several mito-processes, such as fatty acid metabolism, glycolysis or mitochondrial dynamics (Figure 6 b). Remarkably, quantitative proteome data from RPE_T21 c1 cells suggested that all mitochondria-encoded genes involved in OXPHOS, as well as the majority of nuclear-encoded OXPHOS-genes are downregulated (Figure 6 c). In conclusion, mitoXplorer analysis revealed significant differences in mito-gene expression between the different cell lines. Importantly in RPE_T21 cells, proteome data show a remarkable difference to transcriptome data.

**Figure 6:**
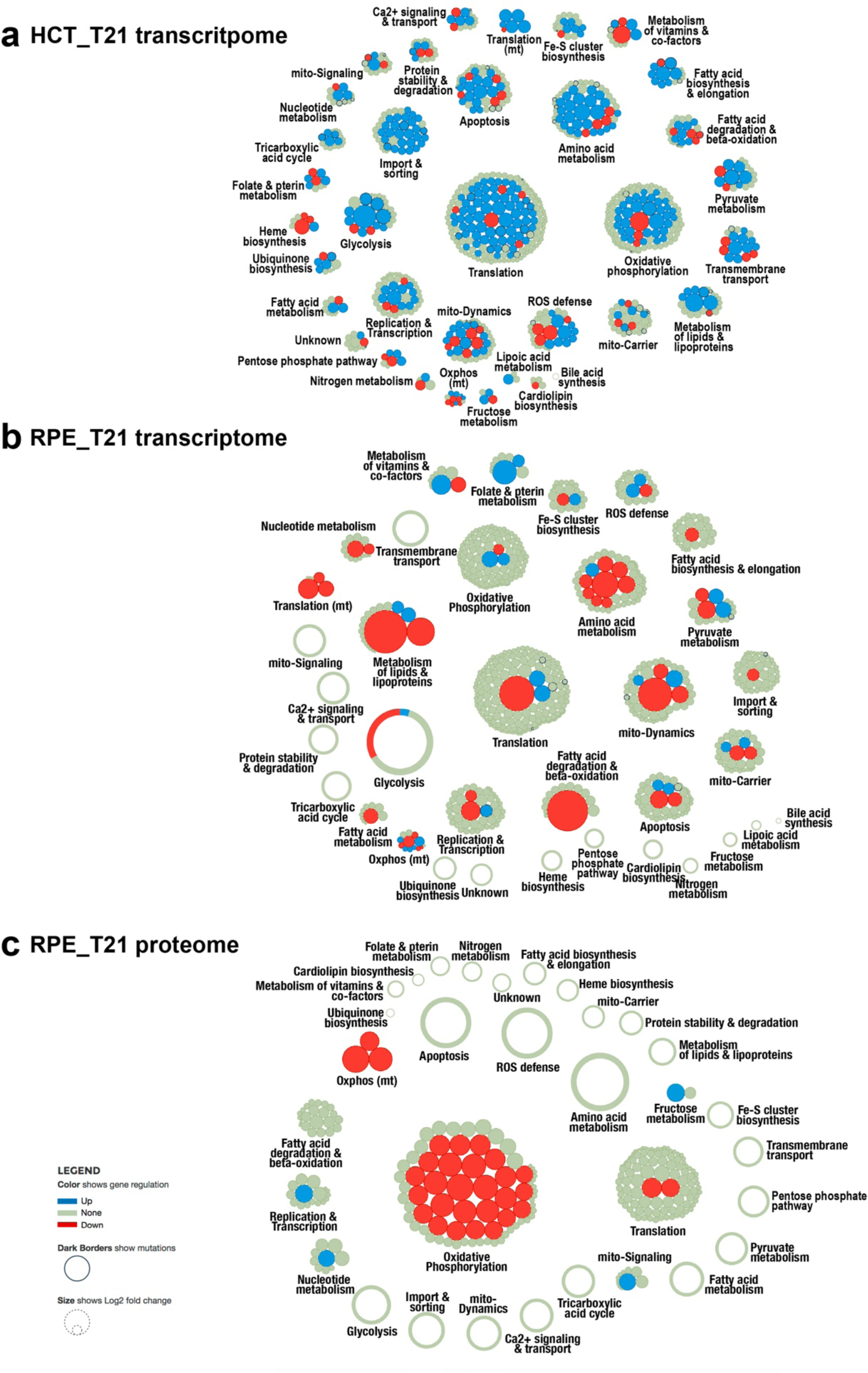
Interactome View of the transcriptome and proteome of cell lines carrying trisomy 21. Trisomy 21 samples were compared against their wild-type counterpart. Transcriptomic analysis of **(a)** HCT116_T21 (trisomy 21 against wild-type) and **(b)** RPE21_T21 (trisomic against wild-type); **(c)** proteomic analysis of RPE_T21 cells (trisomy 21 against wild-type). Transcriptome changes are different between the two trisomy 21 cell lines HCT116 and RPE1. Expression changes of HCT_T21 cells are mild and genes tend to be upregulated **(a)**, while some genes are strongly downregulated in RPE_T21 cells **(b)**. The transcriptome **(b)** and the proteome **(c)** of RPE_T21 cells respond quite differently, with a strong down-regulation of components of the process oxidative phosphorylation (OXPHOS) at proteome level, which is not observed on transcriptome level. Most genes differentially expressed at transcript level, on the other hand, show no significant changes on proteome level. Red bubbles indicate downregulation, blue ones indicate upregulated genes. The size of the bubble corresponds to the log2 fold change.

### mitoXplorer reveals mitochondrial ribosomal assembly defects in RPE_T21 cell lines

To investigate the differences further, we next performed a more detailed analysis of expression changes in these T21 cell lines using Comparative Plots in mitoXplorer. Transcriptome and proteome data from RPE_T21, but not from HCT_T21 cell lines revealed that several subunits of the small mitochondrial ribosome (mitoribosome) were significantly downregulated on either RNA or protein level, or both (Figure 7 a). MRPS21 was strongly reduced on RNA- and protein-level. The genes MRPS33, MRPS14 and MRPS15 were largely normal on RNA level, while their protein levels decreased more than 2-fold (log2FC: MRPS33: −2.147; MRPS14: −1.827; MRPS15: – 1.057). The mitoribosome subunits are encoded in the nuclear genome and their protein products are imported into the mitochondria, where they assemble with mitochondrial ribosomal RNAs to form the large and small subunits of the mitoribosome. The mitoribosome is responsible for translating the 13 mt-mRNAs encoded in the mitochondrial genome, all of which code for key subunits of the respiratory chain required for OXPHOS [80,81]. In accordance with a disrupted mitochondrial translation machinery, all quantifiable mitochondria-encoded OXPHOS proteins (Complex I: MT-ND1 and MT-ND5; Complex IV: MT-CO2) were severely diminished on protein-, but not on RNA-level in RPE1_T21 cells (Figure 7 b,c).

**Figure 7:**
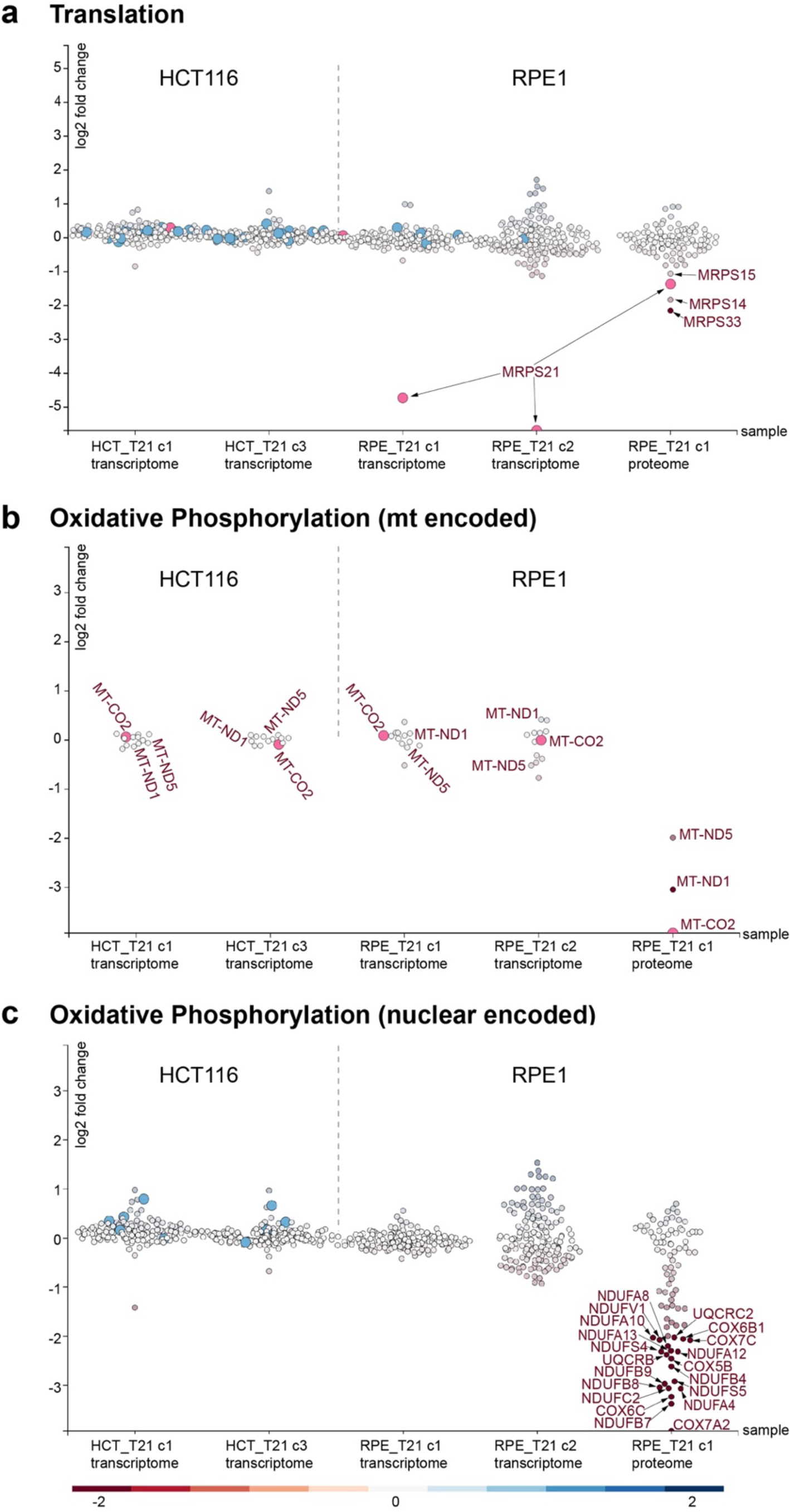
Scatterplots of translation, mitochondrial- as well as nuclear components of oxidative phosphorylation of trisomy 21 cells. **(a)** In the mitochondrial process translation, MRPS21 is strongly down-regulated on transcriptome level in RPE_T21 cells as compared to RPE1 wild-type (wt) cells. No change is observed in HCT_T21 cells. On proteome level, several components of the mitoribosome small subunit (SSU) are down-regulated in RPE_T21 cells. **(b)** Transcript levels of mitochondrial-encoded genes of oxidative phosphorylation (OXPHOS) are not affected. **(c)** A significant number of components of OXPHOS are down-regulated on protein-level in RPE_T21 cells, while no significant or only mild reduction can be observed on transcriptome level in trisomy 21 cell lines. Scatterplots are taken from the mitoXplorer comparative plot interface. Each bubble represents one gene, pink highlighted dots are selected, light blue dots indicate mutated genes. On the y-axis, the log2 fold change is plotted, the cell lines (transcriptome of HCT116 T21 (HCT_T21) clone 1 (c1) and clone 3 (c3) vs wild-type, as well as transcriptome of RPE1 T21 (RPE_T21) clone 1 (c1) and clone 2 (c2) vs wild-type and proteome of RPE1 T21 clone 1 (RPE_T21 c1) vs wild-type) are plotted on the x-axis. The gene highlighted in pink has been selected on the web-server: MRPS21 for the process translation; MT-CO2 for the process mt oxidative phosphorylation; and no gene has been selected in the process oxidative phosphorylation.

Interestingly, 36 of the quantifiable OXPHOS proteins encoded in the nuclear genome were also found to be down-regulated at the proteome, but not at transcriptome level in RPE_T21 cells (Figure 7 c). These include subunits of the NADH dehydrogenase (complex I), ubiquinol-cytochrome c reductase (complex III) and cytochrome c oxidase (complex IV). It is important to note that there is no general down-regulation of mitochondrial proteins in these cells and only a few, specific proteins are strongly downregulated (Figure 6 c). Together, these data demonstrate the power of mitoXplorer to help identify the cause of important changes in mito-gene expression, here the downregulation of mitoribosomal subunits at the transcription level and the resulting cause, in this case the downregulation of the majority of OXPHOS proteins.

### RPE_T21 cells are defective in oxidative phosphorylation

The massive downregulation of OXPHOS proteins in RPE_T21 cells suggests that these cells should suffer from a severe OXPHOS deficiency. To test this hypothesis experimentally, we analyzed cellular respiration and glycolysis in T21 cell lines using a Seahorse XF96 analyzer to quantify oxygen consumption rate (OCR) as an indicator of mitochondrial respiration (Figure 8 a-d, f), as well as the proton production rate (PPR) as an indicator of glycolysis (Figure 8 e, g). In intact RPE_T21 cells, we indeed observed dramatically reduced levels of cellular respiration in comparison to the diploid control (Figure 8 a).

**Figure 8:**
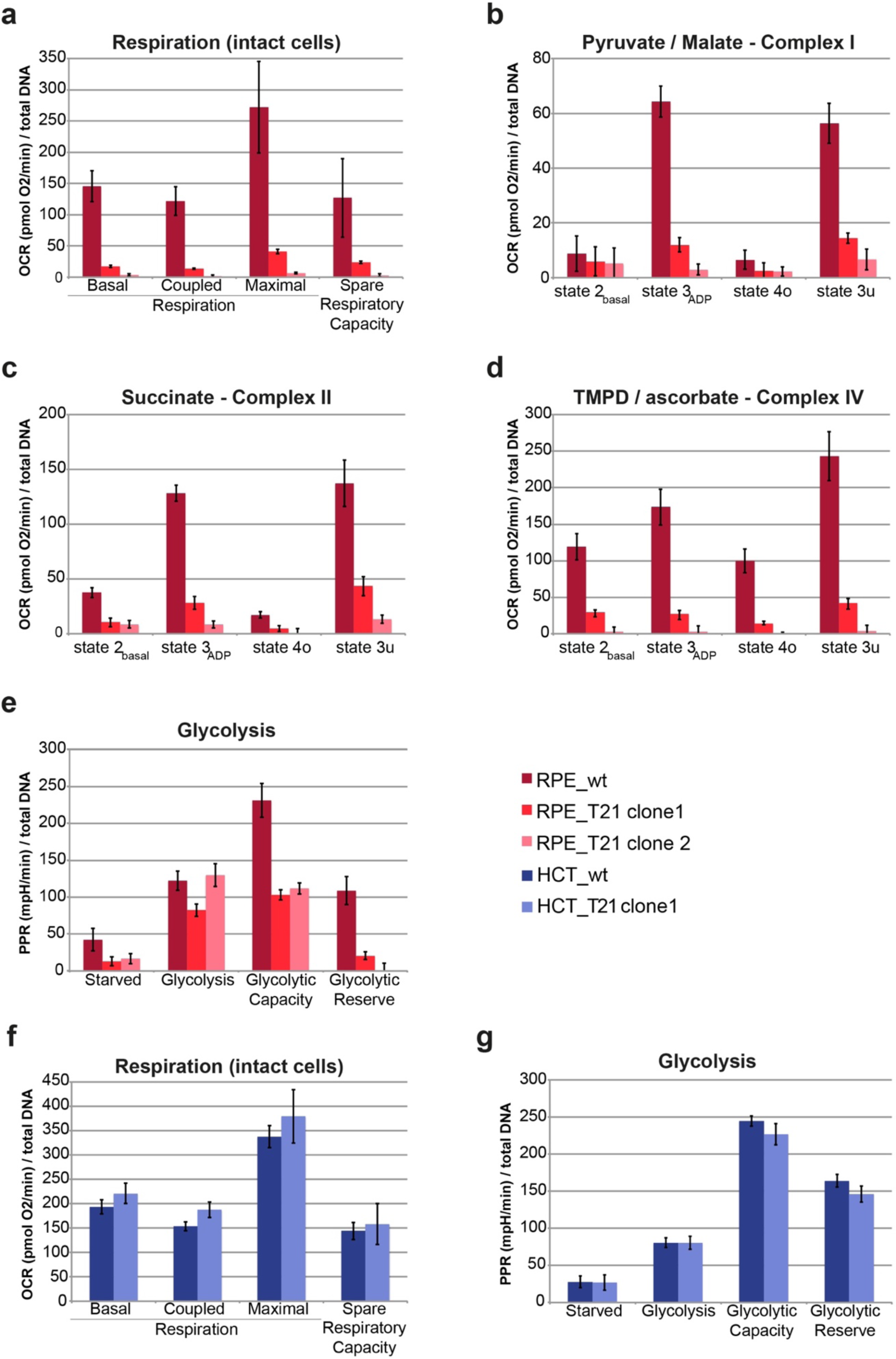
Mitochondrial respiration and glycolysis is strongly affected in RPE_T21 cells and not affected in HCT_T21 cells. **(a)** Respiration in intact RPE_T21 cells is greatly decreased compared to wild-type. **(b – d)** Permeabilized RPE_T21 cells supplemented for substrates of complex I, II and IV as indicated in the header of each plot, showed equally dysfunctional OXPHOS, suggesting a general break-down of the respiratory chain. **(e)** RPE_T21 cells do not have any spare glycolytic reserve. Respiration **(f)**, as well as glycolysis **(g)** is virtually unchanged in HCT_T21 cells compared to their wild-type counterparts. Bright red: RPE_T21 clone 1; light red: RPE_T21 clone 2; dark red: RPE wild-type; dark blue: HCT wild-type; light blue: HCT_T21 clone 1. Measurements of cellular respiration in intact and permeabilized cells, as well as glycolytic potential were done using the Seahorse Bioscience XF Extracellular Flux Analyzer (Seahorse Biosciences). The experiments were performed using the mitochondrial and glycolytic stress test assay protocol as suggested by the manufacturer; the rate of cellular oxidative phosphorylation (oxygen consumption rate (OCR)) and glycolysis (cellular proton production rate (PPR)) were measured simultaneously.

As a complex I deficiency has been reported in trisomy 21 patients [58], we next asked whether RPE_T21 cells selectively suffer from a complex I deficiency, or whether the entire respiratory chain is affected, as suggested by our proteomics data. We used permeabilized cells to test each individual complex with the Seahorse analyzer, supplementing with pyruvate/malate, succinate and TMPD/ascorbate for assessing complex I, II or IV functionality, respectively. As expected from our proteomics analysis, RPE_T21 cells displayed a severe deficiency of the entire respiratory chain (Figure 8 b-d). The glycolytic rate of RPE_T21 cells in the presence of glucose was similar to the diploid control cells. Inhibition of ATP-production was not able to stimulate the cells to a higher glycolytic rate (Figure 8 e), which agrees with the already low OXPHOS levels observed in these cells. HCT_T21 cells, on the other hand, displayed normal respiration, as well as glycolysis (Figure 8 f, g). This suggests that the respiratory chain, as well as the mitochondrial translational machinery is not generally affected in all T21 cells. Taken together, mitoXplorer uncovered OXPHOS deficiencies in RPE_T21 cells, which we verified experimentally, demonstrating the power of an in-depth analysis of mitochondrial expression dynamics to identify the potential molecular cause of the observed phenotype.

### Quantification of mitochondrial network morphology using mitoMorph

We further wanted to investigate, if T21 and the defective OXPHOS had a consequence on mitochondrial morphology and the mitochondrial network structure was changed in T21 cell lines. To quantify mitochondrial morphology in RPE_T21 cells, we stained mitochondria using the MitoTracker Deep Red FM dye. In order to quantify the characteristics of mitochondrial morphology, we developed a new Fiji plug-in for quantification of mitochondrial network features, which we called mitoMorph (Figure 9 a,b). MitoMorph is based on the scripts provided by Leonard, et al. [82] for quantifying mitochondrial network features such as *filaments*, *rods*, *puncta* and *swollen* mitochondrial structures (see Materials and Methods for implementation details). MitoMorph reports the percentages of filaments, rods, puncta and swollen for each individual cell, as well as for all selected cells in a batch analysis. Moreover, it provides the lengths and areas of filaments and rods. Figure 9 (c-f) shows the distribution of mitochondrial network features for the two wild-type and T21 cell lines.

**Figure 9:**
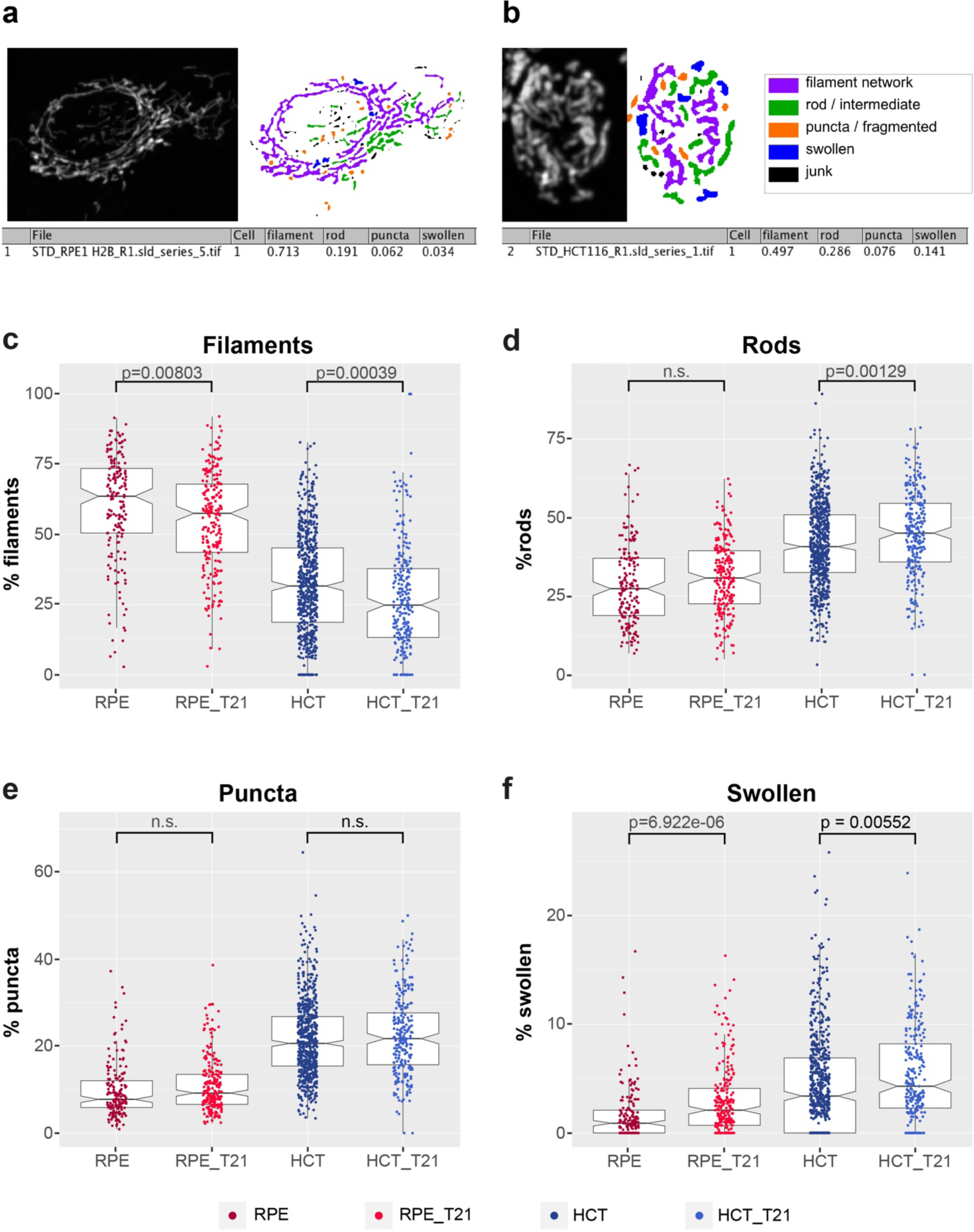
Mitochondrial morphology is slightly changed in trisomy 21 cells. We have stained the mitochondrial network and analyzed the network morphology, measuring the percentage of filaments, rods, puncta and swollen using the Fiji plug-in mitoMorph. **(a)** Sample of mitoMorph analysis of an RPE1 wild-type cell and **(b)** of an HCT116 wild-type cell. Filament networks are highlighted in lilac, rods in green, puncta (referring to fragmented mitochondria) are highlighted in orange and swollen ones in blue. The percentage filaments, rods, puncta and swollen mitochondria are automatically scored and reported to the user. **(c)** the percentage of filaments is slightly, but significantly reduced in RPE_T21, as well as HCT_T21 cells compared to wild-type. **(d)** There is no significant change in the percentage of rods in RPE_T21 cells and slightly higher percentage in HCT_T21 cells compared to wild-type. **(e)** The percentage of puncta is unchanged in both T21 cell lines. **(f)** There are significantly more swollen mitochondria in both, RPE_T21, as well as HCT_T21 cells compared to wild-type. Underlying numerical values are provided in Additional File 4, Supplementary Table S3, sample images are shown in Additional File 1, Supplementary Figure S4.

MitoMorph analysis revealed that in both backgrounds, T21 cells had fewer mitochondrial filaments than their wild-type counterparts, but instead possessed a slightly higher number of rods, which was significant in HCT_T21 cells. Both T21 cell lines had significantly more swollen structures than their wild-type counterparts. Length and area distribution of filaments and rods were not significantly different between the wild-type and the trisomy 21 cells (Additional File 1, Supplementary Figure S2 a-d). In conclusion, mitochondrial morphology based on light-microscopy is mildly affected in trisomy 21.

### Data integration with publicly available trisomy 21 datasets

After discovering this differential OXPHOS defect in our RPE_T21 cell lines, we were interested in the overlap of the mito-transcriptome and -proteome of RPE_T21 cells with data from trisomy 21 patients. We used proteomic and transcriptomic data from a monozygotic twin study discordant for chromosome 21 [41,78]. In agreement with our RPE_T21 data, systematic proteome and proteostasis profiling of fibroblasts from monozygotic twins discordant for T21 revealed a significant, although milder down-regulation of the mitochondrial proteome, including proteins involved in OXPHOS, which is not apparent from transcriptomic analysis of the same cells (see Additional File 1, Supplementary Figure S3 a, b). Taken together, the analysis of both datasets with mitoXplorer suggests a strong post-transcriptional effect leading to reduced expression levels of proteins involved in OXPHOS in trisomy 21.

## Discussion

### The web-based mitoXplorer platform for mito-centric data exploration

MitoXplorer is a practical web tool with an intuitive interface for users who wish to gain insight from -omics data in mitochondrial functions. It is the first tool that takes advantage of the breadth of -omics data available to date to explore expression variability of mito-genes and -processes. It does so by integrating a hand-curated, annotated mitochondrial interactome with -omics data available in public databases or provided by the user.

MitoXplorer has been conceived and implemented as a visual data mining (VDM) platform: by iteratively interacting, visualizing and by allowing manipulation of the graphical display of data, the user can effectively discover complex data to extract knowledge and gain deeper understanding of the data. MitoXplorer provides a set of particularly interactive and flexible visualization tools, with a fine-grained, function- as well as gene-based resolution of the data. Clustering, as well as PCA-analysis help in addition to mine a larger number of -omics data effectively by grouping datasets with similar expression patterns.

VDM-based knowledge discovery is offered by a large number of resources and platforms. However, to the best of our knowledge, no currently available tool allows to explore expression variation of a specific subset of genes in a large number of -omics datasets. It permits users to exploit publicly available transcriptome, proteome or mutation data to study the variation and thus, the adaptability of a defined gene set in different conditions or species. While mitoXplorer offers the exploration of mito-genes, we have designed the platform such that users, who wish to download a local version of mitoXplorer can also upload their own interactome, which can be any gene group of interest. Thus, mitoXplorer can be flexibly adjusted to any user-defined interactome set.

### Cell type-specific de-regulation of mito-genes in trisomy 21

We tested mitoXplorer by expression profiling of mito-genes in T21 cell lines. Mito-genes were strongly deregulated in both trisomic cell types tested, the non-cancerous retinal pigment epithelial cell line RPE1 and the cancer cell line HCT116. Yet, the changes in expression were quite different in the two cell lines. It is not unexpected that mito-genes are differentially expressed in different cell types, reflecting the divergent cellular energy- and metabolic demands [20]. Gene expression is moreover tightly regulated in a cell-type specific manner by regulating transcription, translation and the epigenetic state of the cell. Thus, also divergent and cell-type specific expression changes of mito-genes upon introduction of an extra chromosome is not surprising.

### mitoXplorer revealed divergent de-regulation of mitochondrial transcriptome and proteome in trisomy 21

We found a remarkable difference between transcriptome and proteome levels of mito-genes in RPE_T21 cells. In particular the OXPHOS proteins were strongly down-regulated at protein, but not mRNA level. This can be explained by essential components of the respiratory chain being encoded in the mitochondrial genome and thus requiring a functioning mitochondrial replication system, as well as intact mitochondrial transcription and translation. Thus, there is as strong post-transcriptional regulation of the mitochondrial proteome. In case of the RPE_T21 cell line, the disintegration of the mitoribosome and thus a failure of mitochondrial translation is likely causative for the down-regulation of OXPHOS components on protein-level, possibly by proteolysis, as the essential mitochondrial subunits are not produced and thus complexes cannot assemble. This conclusion is further supported by the fact that we could not observe a significant difference in mitochondrial transcript levels, with some mt-mRNAs even being upregulated; thus, mtDNA -maintenance, -replication as well as mito-transcription seem to be unaffected.

MitoXplorer analysis of previously published data of the mito-proteome of fibroblasts isolated from monozygotic twins discordant for T21 [78] revealed a similar post-transcriptional effect as our T21 model cell lines. Taken together, our data uncovered a significant post-transcriptional regulation of the mitochondrial process OXPHOS in trisomy 21 that could bring new insight into the mechanisms of mitochondrial defects in trisomy 21 patients.

### mitoXplorer identified mitochondrial ribosomal protein S21 (MRPS21) as potentially causative for OXPHOS failure

The most notable difference in RPE_T21 cells compared to wild-type is the 10-12 – fold downregulation of mitochondrial ribosomal protein S21 (MRPS21) on transcript level, as well as the downregulation of Mrps21 protein and other proteins of the small and – to a lesser extend – large mitoribosome subunit. Thus, our data suggest that the integrity of the mitoribosome is compromised, leading to its disintegration and subsequently, the downregulation of mitochondrial proteins of the respiratory chain. Mrps21 is a late-assembly component and lies at the outer rim of the body (or bottom) of the small subunit (SSU) of the mitoribosome. Nevertheless, it interacts with a number of other proteins of the SSU and also directly contacts bases of the 12S rRNA [83,84]. Thus, its absence could destabilize the SSU of the mitoribosome. The two most down-regulated proteins are Mrps33 and Mrps14, both of which directly interact with each other and several other proteins in the SSU and are localized to the head of the SSU. Furthermore, together with another down-regulated component, Mrps15, they are proteins that are incorporated late in the mitoribosome assembly process [84]. This raises the possibility that late-assembly proteins disintegrate more readily from the mitoribosome, leading to their enhanced degradation and thus ribosome malfunction.

Based on promoter analysis using MotifMap [85], potential binding motifs of two transcription factors located on chromosome 21, GABPA and ETS2, can be found in the promoter region of the MRPS21 gene. Gabp*α*, which is also known as nuclear respiratory factor 2, has already been implicated in mitochondrial biogenesis by regulating Tfb1m expression [86]: its depletion in mouse embryonic fibroblasts showed reduced mitochondrial mass, ATP production, oxygen consumption and mito-protein synthesis, but had no effect on mitochondrial morphology, membrane potential or apoptosis. Direct or indirect regulation of mitoribosomal proteins could be another regulatory function of this transcription factor. GABPA is not affected on transcriptome level, but is downregulated on protein-level in RPE_T21 cells. ETS2 on the other hand has so far not been implicated in mitochondrial biogenesis or functional regulation.

## Outlook

MitoXplorer is integrating, clustering and visualizing numerical data resulting from expression studies (transcriptome, proteome), as well as mutation data. Thus, it is currently limited to analyzing mito-genes without offering the ability to explore their embedding in a broader, cellular context and thus to learn about potential regulatory mechanisms of observed expression changes of mito-genes. Therefore, in the next release of mitoXplorer, we plan to fully embed mito-genes within the cellular gene regulatory, as well as signaling network by adding information from epigenetic studies (ChIP-seq, methylation data), as well as from the cellular interactome. We will provide the tools to perform enrichment analysis of observed transcription factors binding in the promoter regions of co-regulated mito-genes; and to explore the regulatory network of mito-genes by offering network analysis methods, such as viPEr [87]. Other analysis methods we will provide include correlation analysis, as well as cross-species data mining. Upon user request, we will also add the mitochondrial interactomes of other species. As mitoXplorer stores the mitochondrial interactomes, as well as associated -omics data in a MySQL database, all technical requirements for extending the functionalities of mitoXplorer are already implemented.

## Conclusions

mitoXplorer is a powerful, web-based visual data mining platform that allows users to in-depth analyze and visualize mutations and expression dynamics of mito-genes and mito-processes by integrating a manually curated mitochondrial interactome with – omics data in various tissues, conditions or species. We used transcriptome and proteome data from cell lines with trisomy 21 to demonstrate the value of mitoXplorer in analyzing in detail the expression dynamics of mito-genes and -processes. We have used mitoXplorer to integrate these data with publicly available datasets of patients with trisomy 21. Using mitoXplorer for data mining, we predicted failure of mitochondrial respiration in one of the trisomy 21 cell lines, which we could verify experimentally. Our results demonstrate the power of a visual data mining platform such as mitoXplorer to explore expression dynamics of a specified mito-gene set in a detailed and focused manner, leading to discovery of underlying molecular mechanisms and providing testable hypotheses for further experimental studies.

## Methods

### Implementation of mitoXplorer

#### Web interface of mitoXplorer (Front-end)

The web interface of mitoXplorer at the *front-end* allows users to access, interact and visualize data from its database, including the interactome and expression/mutation data. The interactive elements and visualizations on mitoXplorer are all built with Javascript, a dynamic programming language that enables interactivity on web pages by manipulating elements through DOM (Document Object Model). DOM is a representation of document, such as HTML, in a tree structure, with each element as a node or an object. Through Javascript and its libraries, visualizations in mitoXplorer can react to users’ action and dynamically change the properties (size, color, coordinates) of web elements and display interactivity. All the visualization components in mitoXplorer described below are modular by design and can be deployed individually or incorporated into web platforms easily.

##### Mitochondrial Interactome (D3 – Data binding and selection)

The visualization of the interactome is created with the implementation of a Javascript library, D3 (d3.js) [88]. D3 (Data-driven documents) is capable of binding data, usually in the form of JSON (Javascript-oriented notation), to the elements of the DOM so that their properties are entirely based on given data. In the interactome, D3 creates an SVG (Scalable Vector Graphic) element for each gene within the DOM in the form of a bubble, with sizes and colors dependent on their Log2FC values. The coordinates of bubbles are also calculated according to the data (e.g. the largest one being at the center) so that the layout of the whole interactome is visually appealing. Upon hovering over any bubble (gene), D3 selects the element and passes additional data bound to that element to the corresponding web element (sidebar) for display.

##### Comparative plot (D3 – Transition and sorting)

The comparative plot combines three interdependent visualizations (scatterplot, bar chart and heatmap) built upon D3. Apart from data-binding and selection, these visualizations exploit the functionality of D3 of transition and sorting through its API. In the scatterplot, genes are displayed as nodes, whose colors and position again depend on the data (log2FC). When another mito-process is selected at the bar chart, D3 updates the data bound to the node and the properties of the nodes are changed. The transition (changes in color and position) is smooth and gives users the impression that the visualization is truly dynamic and interactive. D3 can manipulate, not only the elements, but also the data bound to the elements. Upon clicking the dataset or gene names on the heatmap, the data can be sorted accordingly and an index is assigned to each element (tile on the heatmap) to indicate its position.

##### Hierarchical clustering (mpld3 – Visualization in Python implemented in D3)

The heatmap displaying the results of hierarchical clustering is built with mpld3, a Python library that exports graphics made with Python’s Matplotlib-based libraries to JSON objects that could be displayed on web browsers. Mpld3 benefits from D3’s data-binding property and allows users to create a plugin that interacts with the data on the visualization. The advantage of using mpld3 is that the analysis and visualizations made in Python can be directly translated to JSON and deployed in Javascript on webpages without re-programming. In the case of hierarchical clustering, since libraries for both clustering analysis and visualization of results in a heatmap with a dendrogram are available in Python (described below), it is exported to JSON with mpld3, and a Javascript tooltip plugin that allows users to select data or display information with D3.

##### Principle Component Analysis (three.js – 3D visualization)

The visualization of the result of Principle Component Analysis (PCA) is 3-dimensional, with each dimension representing one of the first three Principle Components (PCs). This is achieved through the implementation of three.js, a Javascript library that enables animated 3D graphics to be created and displayed in a web browser. It starts with building a “scene”, or a canvas, on which 3D objects will be created. Then a “camera” is set up that controls the view of objects on the scene from the users’ perspective, such as the field of view (width, height, depth) and its ratio; and a “renderer” that renders the scene at short time intervals so objects are displayed as animated object (either they are animated by themselves or moved around on the scene by users). Objects of different texture, geometry and color, can now be added to and rendered on the scene. Finally, the scene with objects is attached to the DOM of a webpage to become visible. In the PCA visualization, each dataset is represented and rendered as a small sphere, with coordinates (x, y, z) depending on the values of its first three PCs, and colors on the grouping of that dataset. When users drag around on the canvas or zoom in or out, all objects are re-rendered in such a way that the scene appears to be a 3-dimensional space.

#### MitoXplorer Database (*back-end*)

A MySQL database hosted at the *back-end* of mitoXplorer contains the interactomes of mito-genes, including the mito-process, gene ontology and the interactions between gene products; and the expression and mutation data from public databases. Each entry of the expression and mutation data has a foreign link to the interactome and file directory (dataset table). This ensures that the expression and mutation data will be updated together with the interactome, or when a dataset is updated or deleted. Users can upload their own differential expression and/or mutation data, which will be processed and integrated with the interactome by extracting mito-genes, and stored in the mitoXplorer database for up to 7 days.

### Data analysis and communication between front- and back-end

A Python application serves as a bridge between the *front*- and *back-end* of mitoXplorer. Upon the users’ request to access the database or perform analysis at the web interface, an AJAX-asynchronous call directed to the Python application is made, so the request can be performed in the background and the webpage is updated without reloading. The Python application then processes the request by connecting to the MySQL database and analyzes the data retrieved from it. The application also handles the user uploads (e.g. data cleaning) before saving it to the MySQL database. The main libraries used by the Python application for analysis include: 1) Scikit-learn: a machine learning library that provides tools for PCA, to perform dimensionality reduction on the expression of all mito-genes and of each mito-process. The first three principal components are extracted for each dataset. 2) SciPy: a mathematical library that provides modules for Hierarchical Clustering, to calculate 2-dimensional distance matrices between genes and between datasets based on expression values, for each mito-process. 3) Seaborn: a statistical visualization library built on top of SciPy to create heatmaps from the results. All the results are produced in JSON format, which are then sent via the HTTP protocol back to the *front-end* and visualized with Javascript.

The usage of mitoXplorer does not require installation or programming knowledge. Documentation and tutorials are available online and on GitLab (https://gitlab.com/habermannlab/mitox). MitoXplorer is also available for download and installation on a local server, if users wish to build their own gene list and apply the interactive features and database of mitoXplorer, which stores the available expression and mutation data for all genes. Setup instructions are also available on GitLab (https://gitlab.com/habermannlab/mitox). We also provide a docker version of mitoxplorer at (https://gitlab.com/habermannlab/mitox, branch docker-version).

### Transcriptomics and proteomics of aneuploid cell lines

The proteome analysis of the trisomic cell lines was previously described [44,45].

The raw reads from RNA-sequencing were processed to remove low quality reads and adapter sequences, and aligned to the reference genome (hg19) with TopHat2 [89]. The Cufflinks package [90] was used to calculate the expression difference between two samples (aneuploid vs diploid) of multiple replicates and test the statistical significance. Transcriptome and proteome information are available in public repositories: NGS data have been deposited in NCBI’s Gene Expression Omnibus and are accessible through GEO Series accession number GSE102855 and GSE131249. Data from Kühl et al. [46], as well as Liu, et al. [78], Letourneau et al. [41] and Sullivan [42] were uploaded as provided by the authors.

### Data processing of public data and correlational analysis

The public NGS datasets on mitoXplorer were downloaded from GEO; only RNA-seq data, not microarray data are currently uploaded on mitoXplorer. The pre-analyzed data were downloaded and transformed to transcript per million (TPM). Log2FC were calculated for each disease-sample, using the corresponding diploid samples as control (or the mean of normal samples if there were no paired samples). Metadata of the samples (e.g. cell types) was also downloaded and stored in the mitoXplorer database. The links to the experiments for each dataset are available at the DATABASE summary page of mitoXplorer. TCGA differential expression data were downloaded from the NCI GDC Data Portal (https://portal.gdc.cancer.gov/). For calculating differential expression, the log2FC was calculated from TPM (transcripts per million) for each paired sample.

### Cell culture and treatment

The human cell line RPE-1 hTERT (referred to as RPE) was a kind gift by Stephen Taylor (University of Manchester, UK). Human HCT116 cells (referred to as HCT) were obtained from ATCC (No. CCL-247). Trisomic cell lines were generated by microcell-mediated chromosome transfer as described previously [45]. The A9 donor mouse cell lines were purchased from the Health Science Research Resources Bank (HSRRB), Osaka 590-0535, Japan. All cell lines were maintained at 37°C with 5% CO2 atmosphere in Dulbeccós Modified Eagle Medium (DMEM) containing 10% fetal bovine serum (FBS), 100 U penicillin and 100 U streptomycin.

### MitoTracker staining and imaging

Mitochondria were stained in 96-well plates. The cells were incubated for 30 min at 37°C with 100 nM MitoTracker deep Red FM (M22426, Invitrogen ®) dye prior to fixation. Cells were fixed with 3% PFA in DMEM for 5 min at room temperature. After washing twice, 1xPBST, plates were stored with 1xPBS containing with 0.01% sodium azide was added. Plates were stored at 4°C in the dark. Imaging was carried out on an inverted Zeiss Observer.Z1 microscope with a spinning disc and 473 nm, 561 nm and 660 nm argon laser lines. Imaging devices were controlled, captured, stored and processed with the SlideBook Software in Fiji [91]. The images were captured automatically on multiple focal planes (step size 700 nm) with a 40x magnification air objective.

### Metabolic profiling of wild-type and T21 cell lines

RPE and HCT cells and their T21 derivatives were seeded at 25,000 or 36,000 cells/well respectively, on XF96 cell plates (Seahorse Bioscience, Agilent Technologies), 30 hours before being assayed. Optimization of reagents as well as CCCP and digitonin titrations were performed as described by the manufacturer’s protocols (Seahorse Bioscience). The experiments were performed using the mitochondrial and glycolytic stress test assay protocol as suggested by the manufacturer (Seahorse Bioscience, Agilent Technologies). By employing the Seahorse Bioscience XF Extracellular Flux Analyzer, the rate of cellular oxidative phosphorylation (oxygen consumption rate (OCR)) and glycolysis (cellular proton production rate (PPR)) were measured simultaneously.

For OCR measurement, DMEM media was supplemented with 25 mM glucose, 1 mM pyruvate and 2 mM glutamine. Basal rate was recorded and additions for the mito stress test were as follows: 1.5 µM oligomycin, CCCP, 2µM rotenone + 4 µM antimycin A. For PPR measurement, DMEM media was supplemented with 2 mM glutamine. Basal rate was recorded and additions for the glycolysis stress test were as follows: 10 mM glucose, 1.5 µM oligomycin and 100 mM 2-deoxyglucose.

For intact cells, the CCCP concentrations were 7 and 1.5 µM for RPE1 and HCT116 cells, respectively. The assays of intact cells were performed in 96-well plates with at least 10 replicates per cell line. For the permeabilized RPE1 cell lines, the CCCP and digitonin concentrations were 10 µM and 40 µM, respectively. In the case of permeabilized HCT116 cell lines, the CCCP and digitonin concentrations were 12 and 50µM, respectively. For OCR measurement, Mannitol-sucrose buffer (MAS) was prepared according to Seahorse Biosciences. For permeabilization, digitionin was added to MAS buffer together with the respective respiratory substrates: 10mM pyruvate / 2 mM malate, 10 mM succinate / 2 µM rotenone or 0.5 mM TMPD / 2 mM ascorbate / 2 µM antimycin A. Basal respiration was recorded, as were additions of 4mM ADP, 1.5 µM oligomycin, CCCP and 2 uM rotenone ± 4uM antimycin A or 20 mM Na-azide. The assays in permeabilized cells were performed in poly-D-lysine-coated 96-well plates with at least 5 replicates per cell line.

Normalization was performed with the CyQuant cell proliferation assay kit (Life Technologies) in the same plate used for the assay of intact cells; and in a parallel plate for the permeabilized cells. Data analysis was done according to [92].

### The mitoMorph plug-in for morphological characterization of mitochondria by image analysis

Classification and measurement of mitochondria were performed using the software ImageJ [93], complemented with all the default plugins provided by Fiji [91] and with the additional plugin FeatureJ. A set of functions were developed to assist the user in the preparation and analysis of the data, either in interactive or batch processing mode.

Using this toolset, after all the cells of interest were manually outlined in each image, the mitochondria were segmented and characterized. For each processing step, the algorithms used are reported as described in ImageJ, and their parameters are specified in physical units.

The images were pre-processed by first suppressing the background signal (rolling ball background subtraction, kernel radius: 2.5 µm) and then enhancing the mitochondria signal (Laplacian of Gaussian, smoothing scale: 1 µm, followed by contrast limited adaptive histogram equalization, CLAHE, kernel size: 2.5 µm). Mitochondria candidates were obtained by segmentation, using Yen thresholding algorithm [94], and subjected to classification based on a set of determined features.

Objects that were too small were excluded from the analysis, and the remaining ones were assigned to one of four categories: networked, puncta, rods and swollen [82]. Objects that were quasi-round, compact in intensity, and larger than the puncta were classified as swollen. All objects with an intermediate phenotype between fragmented puncta and network of filaments were classified as rods.

Classification was performed by sequentially verifying different selection criteria, one set for each class, based on the following measured features: area (A), aspect ratio (AR), circularity (C), solidity (S), minimum Feret diameter (here indicated as minimum linear extension, MLE) and longest shortest-path (here indicated as extension, E). While all the other measures are directly derived from the segmentation, the extension is measured as the longest shortest-path between any two end points in the skeleton derived from the segmentation. The selection criteria are evaluated sequentially as reported in Additional File 4, Supplementary Table S3.

We would like to note that analysis of mitochondrial morphology on projected images is limited, as mitochondrial structures might not be resolved properly.

### Image analysis using mitoMorph and data processing

Image processing and analysis was done in Fiji. Image stacks were Z-projected, cells were manually selected and the resulting images were saved for further batch processing using mitoMorph. Resulting network statistics of mitochondrial features for each individual cell were used for further processing (Additional File 4, Supplementary Table S3). All statistical processing and data visualization of mitoMorph results was done using R.

## Declarations

### Acknowledgements

We want to thank Michael Volkmer for help and advice in web-server management and development. We want to thank Stephen Taylor (University of Manchester, UK) for providing cell lines. We thank Alice Carrier, Friedhelm Pfeiffer and Frank Schnorrer for critical reading of the manuscript. We thank the Max Planck Society, the Max-Planck Institute for Biochemistry, the Aix-Marseille University, the CNRS and the Institute of Developmental Biology Marseille (IBDM) for their support.

### Availability of data and materials

The mitoXplorer web-server is freely available at http://mitoxplorer.ibdm.univ-mrs.fr/. The source code of mitoXplorer is available at https://gitlab.com/habermannlab/mitox. The pipeline for differential expression analysis and mutation calling of RNA-seq data is available at https://gitlab.com/habermannlab/mitox_rnaseq_pipeline. MitoMorph is freely available at https://github.com/giocard/mitoMorph. RNA-seq data published with this study are available via the Gene Expression Omnibus (GEO) database (accession numbers: GSE131249).

### Details on software provided in this manuscript

Project name: mitoXplorer

Project home page: http://mitoxplorer.ibdm.univ-mrs.fr/

Archived version: https://gitlab.com/habermannlab/mitox

Operating system(s): Platform independent

Programming language: JavaScript, PHP, MySQL, Python

Other requirements: none

License: GNU public license

Any restrictions to use by non-academics: none

Project name: mitoMorph

Project home page: https://github.com/giocard/mitoMorph

Archived version: https://github.com/giocard/mitoMorph

Operating system(s): Platform independent

Programming language: Groovy, ImageJ Macro

Other requirements: ImageJ/Fiji software

License: GNU Public License

Any restrictions to use by non-academics: none

### Author’s contributions

AY and PK were the main developers of the mitoXplorer web-server with the help of JV, SG and AB. JV developed the interactome view of the web-server. Data analysis was done by AY, PK, MD and BHH. MitoTracker staining and imaging was carried out by MD, CM and FP carried out metabolic measurements. Handling of cells and cell culture was done by MD and ZS. GC conceived and developed the mitoMorph Fiji plugin, image analysis with mitoMorph was done by BHH. The project was conceived by BHH, the manuscript was written by AY and BHH with contributions from SZ, CG, MD and CM. All authors read and approved the final version of the manuscript.

### Ethics approval and consent to participate

Not applicable.

### Additional files

Additional file 1: Figures S1-S4. Supplementary figures. (PDF format).

Additional file 2: Supplementary table S1 a-c. (Excel format).

Additional file 3: Supplementary table S2 a-e. (Excel format).

Additional File 4: Supplementary table S3 a-d. (Excel format).

## Additional file 1 – Supplementary Figures

**Supplementary Figure S1:**
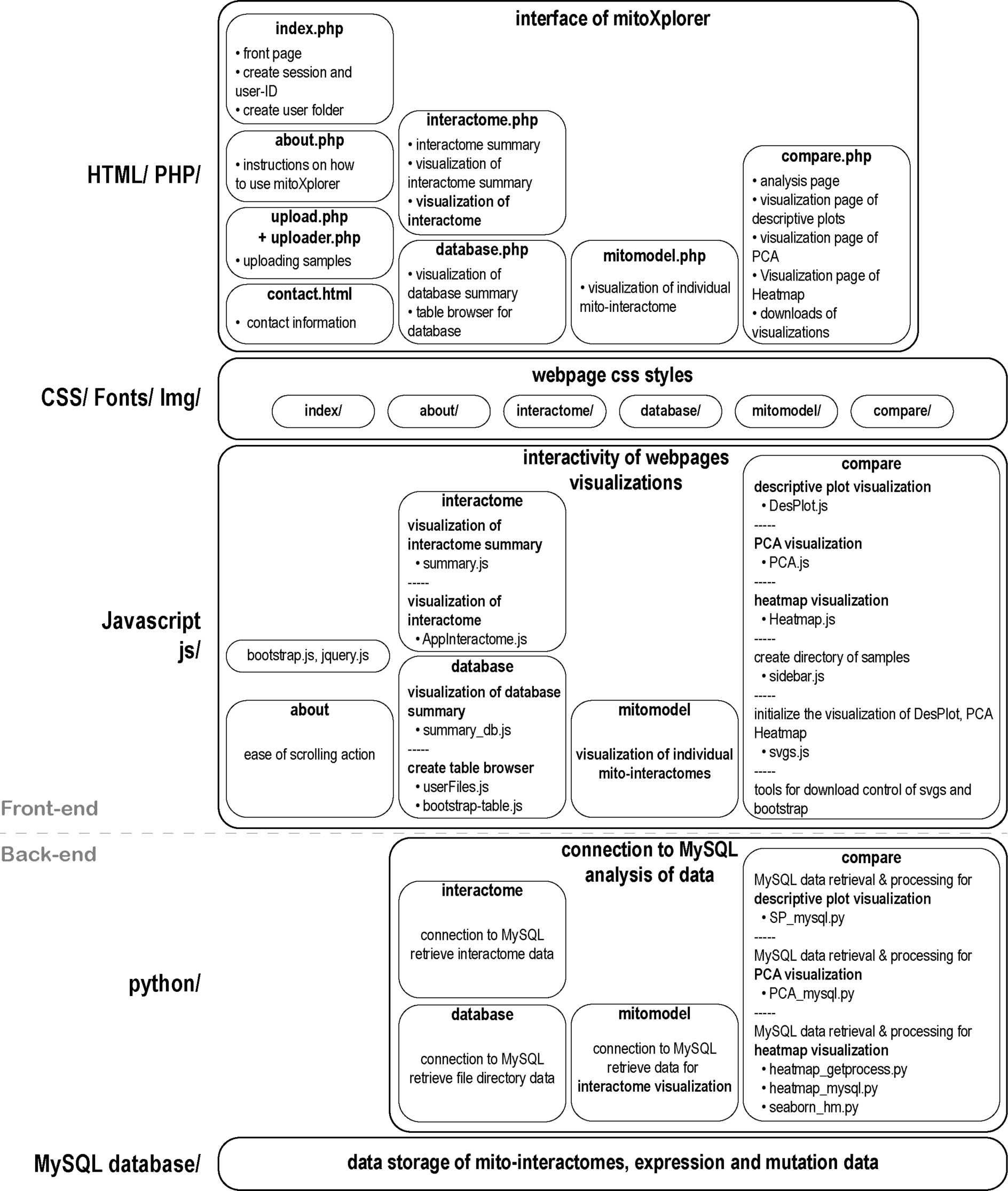
programmatic skeleton of the mitoXplorer web-platform. In the *back-end*, A MySQL database stores the mito-interactomes, as well as expression and mutation data that are publicly available. User-uploaded data are stored temporarily and only available to the user. A set of python-scripts connect to the MySQL database for data retrieval of both, mito-interactomes and expression and mutation data. The mitomodel script connects to the MySQL database directly for the visualization of the Interactome View. A set of scripts perform comparative analysis, for generating Comparative Plots, Heatmap and PCA visualization. In the *front-end*, a set of javascripts handle the visualizations of the plots: the ‘interactome’ and ‘database’ scripts handle the data presentation of the mito-interactome and the available public data for the web-site; mitomodel visualizes the Interactome View and the scripts in the compare box are responsible for visualizing Comparative Plot, Heatmap and PCA. The CSS layer handles the css-styles of the page and finally, the HTML/PHP layer creates the actual interface for the user.

**Supplementary Figure S2:**
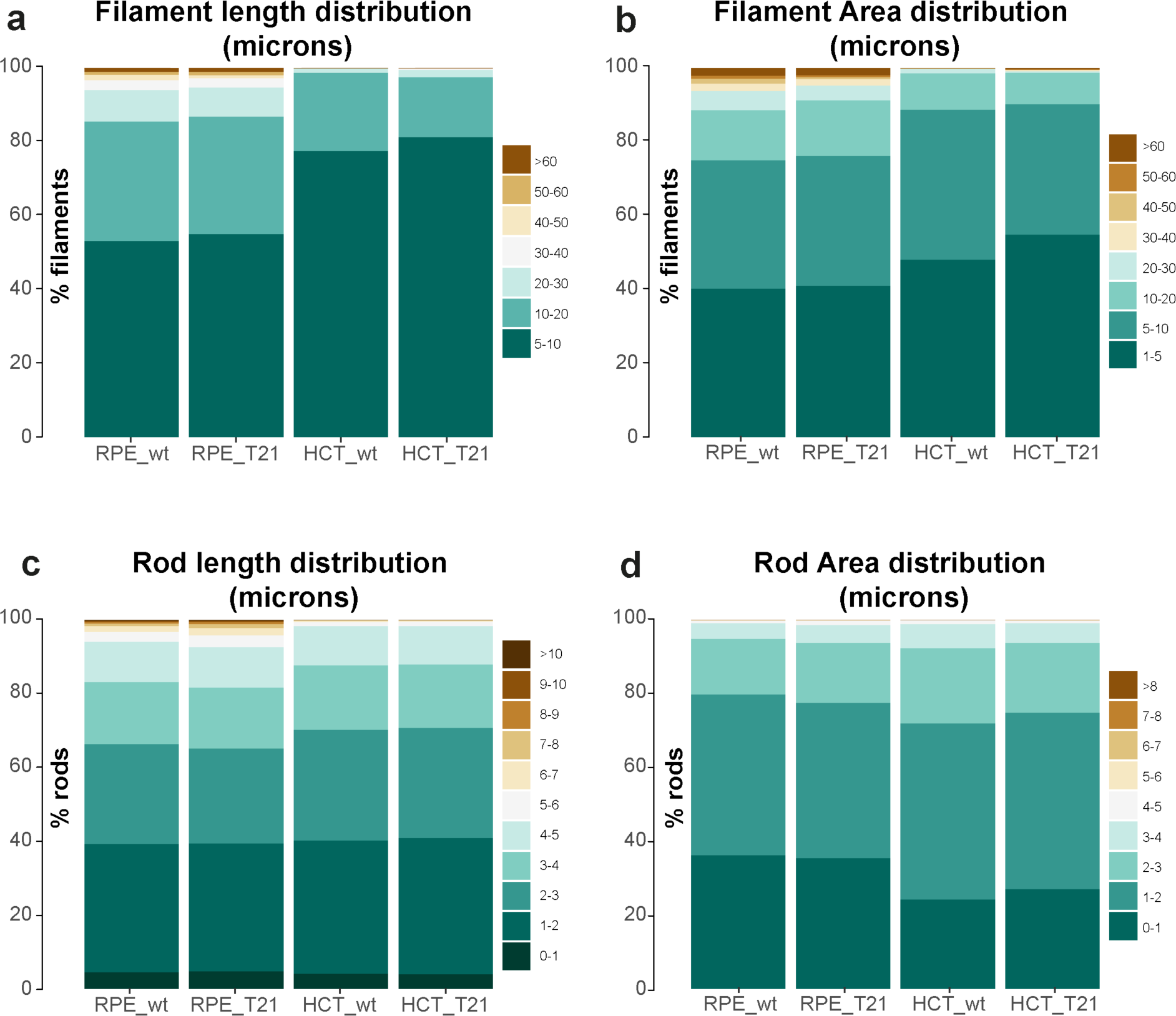
Length and area distribution of filaments and rods in wild-type and T21 derived RPE1 and HCT116 cells. **(a)** stacked bar-plots of filament length distribution of RPE1 wild-type (labeled RPE_wt), RPE1 21/3 (labeled RPE_T21), HCT116 wild-type (labeled HCT_wt) and HCT116 21/3 (labeled HCT_T21) cells. Overall, shorter filaments are more frequent in HCT116 than in RPE1 cells. In T21, filaments tend to be slightly shorter. **(b)** stacked bar-plots of filament area distribution of RPE_wt, RPE_T21, HCT_wt wild-type and HCT_T21 cells. Overall, less area is occupied by filaments in HCT116 than in RPE1 cells. In HCT_T21 cells, a notably smaller area is assigned to filaments, while in RPE_T21 cells, this change is much less pronounced. **(c)** stacked bar-plots of rod length distribution of RPE_wt, RPE_T21, HCT_wt and HCT_T21 cells. Overall, in the range between 4 and 10 microns, more rods are found in RPE1 cells. Between wild-type and T21 cells, no real length difference is observable. **(d)** stacked bar-plots of rod area distribution of RPE_wt, RPE_T21, HCT_wt and HCT_T21 cells. Overall, there is a tendency of slightly larger rod areas in HCT116 cells. In HCT116 cells, rods seem to occupy slightly smaller areas when carrying the extra copy of chromosome 21.

**Supplementary Figure S3:**
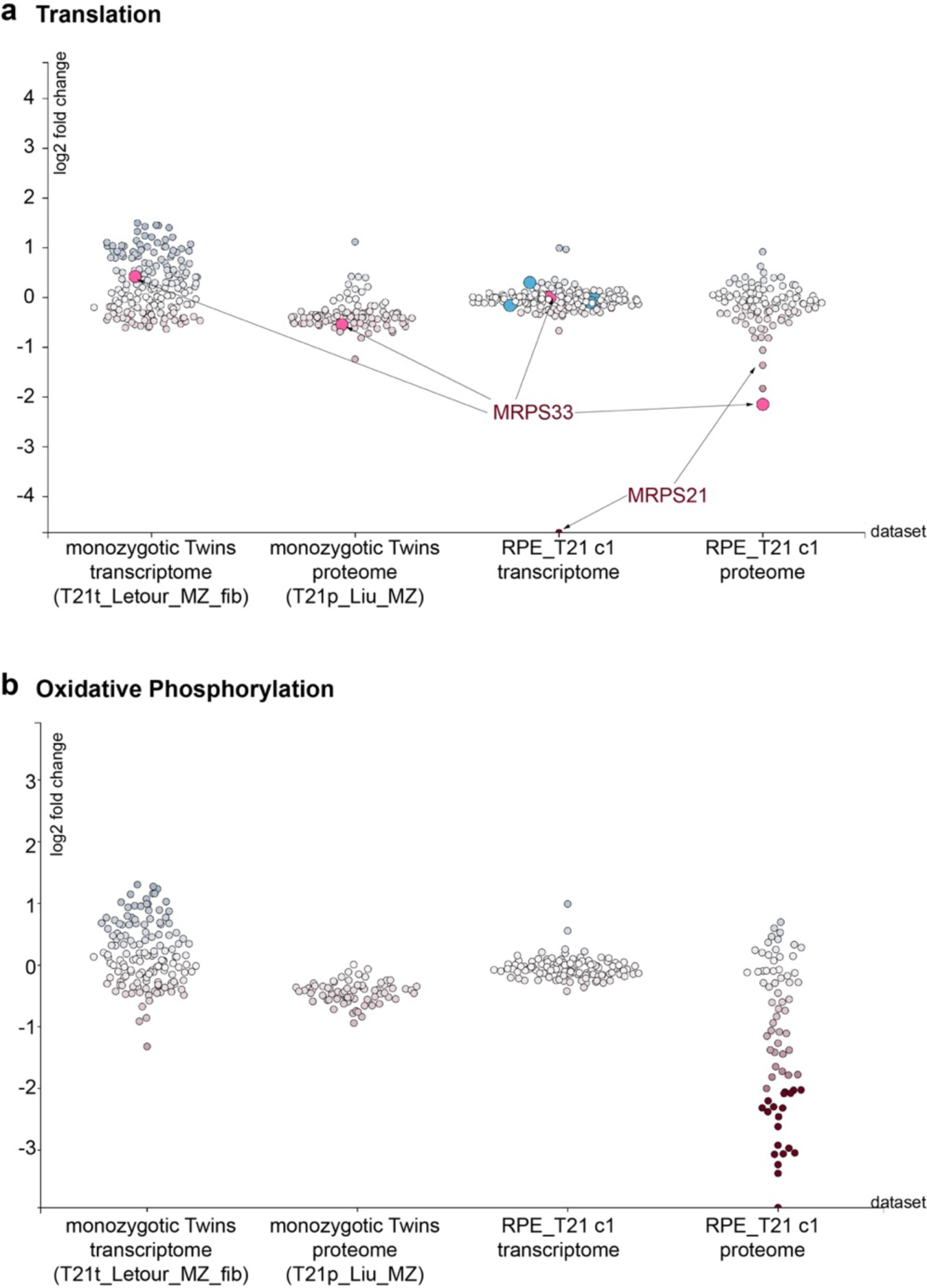
mitoXplorer scatterplot of Translation and nuclear-encoded Oxidative Phosphorylation of fibroblasts of monozygotic twins discordant for trisomy 21 (T21_MZ) and RPE1 T21 cells. **(a)** The mRNA of mitoribosome small subunit component MRPS21 is strongly down-regulated only in RPE_T21 cells and is mostly unaffected in monozygotic twins discordant for T21 (T21_MZ fibroblasts: T21_Letour_MZ_fib, T21_Liu_MZ). Mitoribosome proteins are significantly downregulated in RPE_T21 cells and mildly affected in T21_MZ fibroblasts. **(b)** Oxidative Phosphorylation components encoded in the nucleus are downregulated on protein level in both, RPE_T21, as well as T21_MZ fibroblasts, whereby deregulation is milder in T21_MZ. In both conditions, the Oxidative Phosphorylation transcriptome is mostly unaffected.

**Supplementary Figure S4:**
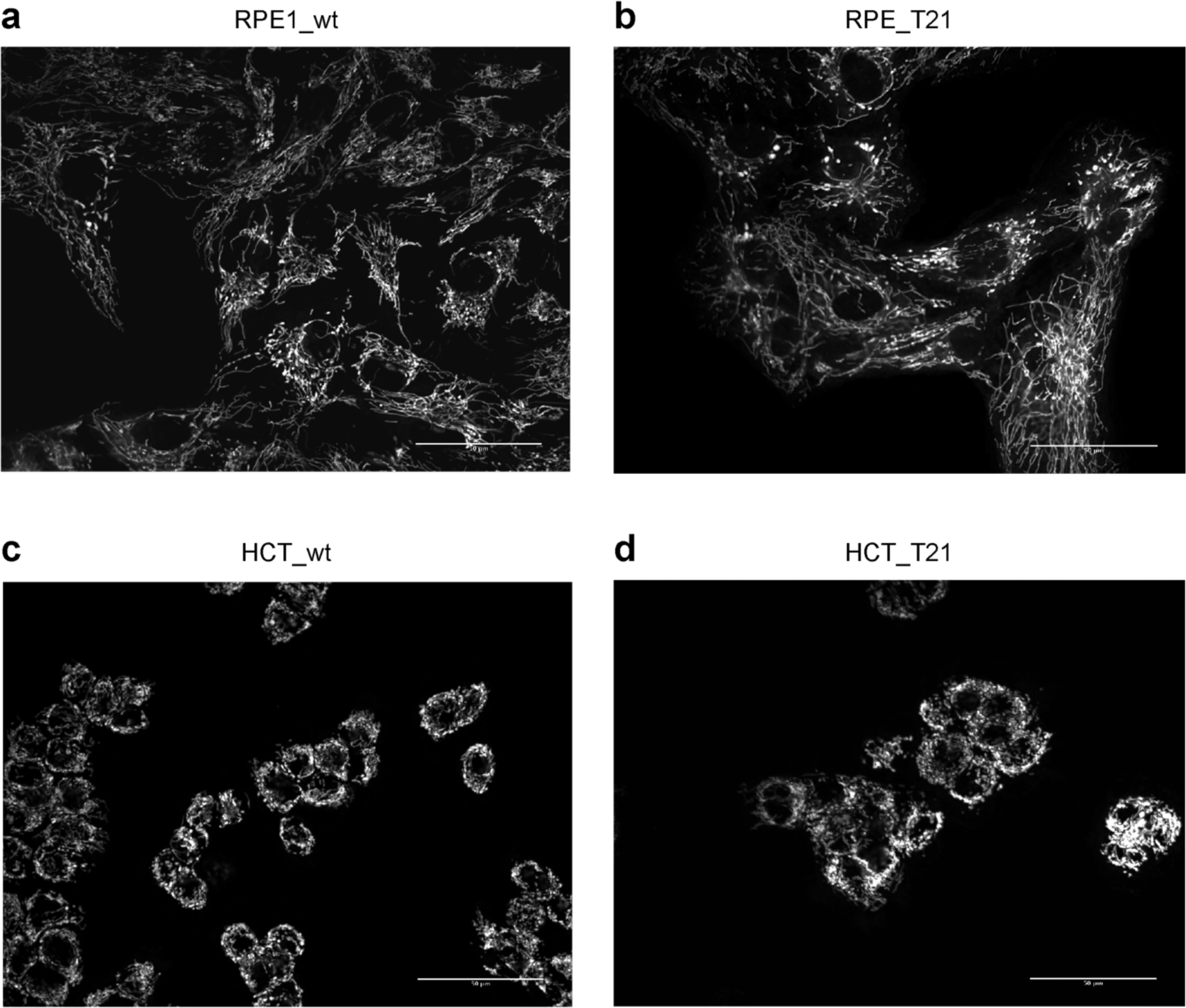
Mitochondrial network and mitoMorph based image analysis of wild-type and T21 cells. MitoTracker stainings **(a-d)** of RPE_wt **(a)** and RPE_T21 **(b)**, as well as HCT_wt **(c)** and HCT_T21 **(d)**. **(a, b)** The mitochondrial network is largely intact in RPE_T21 cells, with only slightly lower percentage filaments and an increased number of swollen mitochondria. **(c, d)** In HCT116 cells, the mitochondrial network is overall less abundant, with more rod-like and fragmented mitochondria (puncta). With trisomy 21, cells show an even more pronounced presence of rods at the cost of longer filaments, as well as more puncta and swollen mitochondria. The scale bar is 50 μm. Mitochondria were stained with MitoTracker deep Red FM from Invitrogen. Staining was done in 96-well plates. The cells were incubated for 30 min at 30°C with 100 nM MitoTracker dye prior to fixation. Cells were fixed with 3% PFA in DMEM for 5 min at room temperature. After washing with 1xPBS, 1xPBS with 0.02% sodium azide was added. Plates were stored at 4°C in the dark. Imaging was carried out on an inverted Zeiss Observer.Z1 microscope with a spinning disc and 473 nm, 561 nm and 660 nm argon laser lines. The images were captured automatically on multiple focal planes (step size 700 nm) with a 40x magnification air objective. Image stacks were Z-projected using Fiji for further analysis.

